# Chromatin endogenous cleavage provides a global view of yeast RNA polymerase II transcription kinetics

**DOI:** 10.1101/2024.07.08.602535

**Authors:** Jake VanBelzen, Bennet Sakelaris, Donna Garvey Brickner, Nikita Marcou, Hermann Riecke, Niall Mangan, Jason H. Brickner

## Abstract

Chromatin immunoprecipitation (ChIP-seq) is the most common approach to observe global binding of proteins to DNA *in vivo*. The occupancy of transcription factors (TFs) from ChIP-seq agrees well with an alternative method, chromatin endogenous cleavage (ChEC-seq2). However, ChIP-seq and ChEC-seq2 reveal strikingly different patterns of enrichment of yeast RNA polymerase II. We hypothesized that this reflects distinct populations of RNAPII, some of which are captured by ChIP-seq and some of which are captured by ChEC-seq2. RNAPII association with enhancers and promoters - predicted from biochemical studies - is detected well by ChEC-seq2 but not by ChIP-seq. Enhancer/promoter bound RNAPII correlates with transcription levels and matches predicted occupancy based on published rates of enhancer recruitment, preinitiation assembly, initiation, elongation and termination. The occupancy from ChEC-seq2 allowed us to develop a stochastic model for global kinetics of RNAPII transcription which captured both the ChEC-seq2 data and changes upon chemical-genetic perturbations to transcription. Finally, RNAPII ChEC-seq2 and kinetic modeling suggests that a mutation in the Gcn4 transcription factor that blocks interaction with the NPC destabilizes promoter-associated RNAPII without altering its recruitment to the enhancer.

## Introduction

In eukaryotes, differential expression of the genome is achieved primarily through regulated RNA polymerase II (RNAPII) transcription. Since its discovery (Roeder and Rutter, 1969), transcription by RNAPII has been the focus of intense study using a variety of methods. From biochemical, structural and genetic studies, a consensus has emerged for the mechanism of RNAPII transcription (Figure 1; Schier and Taatjes, 2020). For genes that are dependent on enhancers, sequence-specific transcription factors (ssTFs) bind to enhancers and recruit co-activators like histone acetyltransferases and chromatin remodelers as well as Mediator (Fishburn et al., 2005; Green, 2005; Prochasson et al., 2003; Ptashne and Gann, 1997). Co-activators facilitate the removal of nucleosomes from the promoter, allowing binding of TFIID (TATA binding protein), which recruits additional general transcription factors (GTFs; TFIIA, TFIIB, TFIIF) and ultimately RNAPII (Figure 1). Last, TFIIE and TFIIH are recruited to complete the formation of the preinitiation complex (PIC). Through Mediator, ssTFs interact with RNAPII to stabilize the PIC (Abdella et al., 2021; Richter et al., 2022). TFIIH stimulates initiation by both unwinding the DNA and by phosphorylating the RNAPII carboxyl terminal domain on Serine 5 (Figure 1, inset; Cadena and Dahmus, 1987; P. Komarnitsky et al., 2000; Lu et al., 1991). In metazoans, regulatory factors (negative elongation factor and DRB-sensitive factor, DSIF) cause RNAPII to pause after initiation, leading to an accumulation of RNAPII downstream of the transcriptional start site (Adelman and Lis, 2012; Core and Adelman, 2019). The P-TEF-b kinase releases RNAPII from pausing by phosphorylation of these factors and RNAPII on Serine 2, leading to elongation (Marshall and Price, 1995). Finally, transcription of a polyadenylation sequence both causes RNAPII to pause, simulating cleavage and polyadenylation (Figure 1; Nag et al., 2007; Orozco et al., 2002).

**Figure 1.**
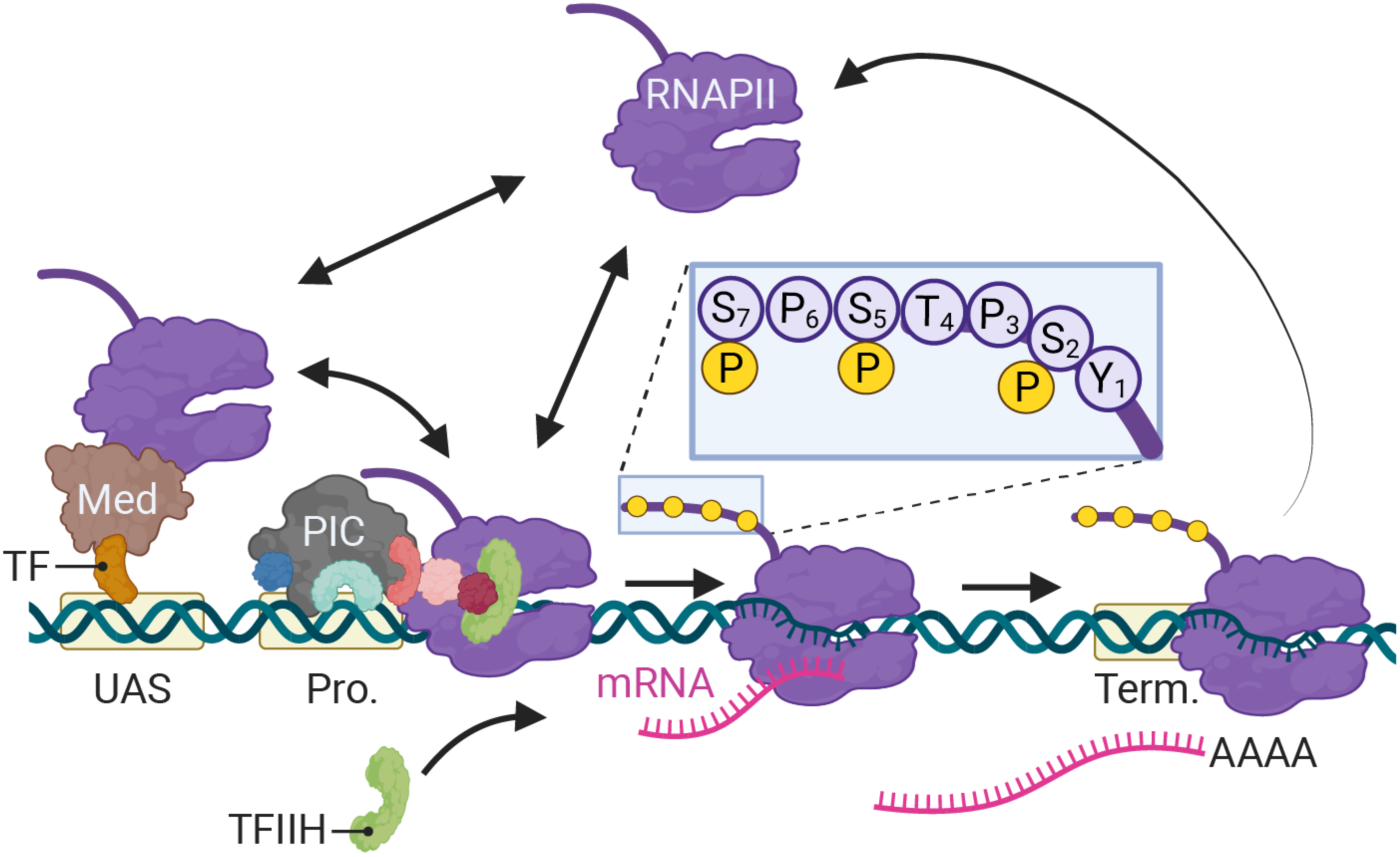
Schematic of RNAPII-mediated transcription in *S. cerevisiae*. Two alternative mechanisms of RNAPII recruitment are shown: 1) direct recruitment to the promoter and 2) recruitment to the UASs facilitated by a ssTFs and coactivators such as Mediator, followed by transfer to the promoter. After RNAPII associates with the promoter, TFIIH is recruited, leading to phosphorylation of Serine 5 (inset) in the carboxyl terminal domain by the TFIIH-associated kinase Kin28 and initiation. RNAPII elongation through the transcribed region is associated with phosphorylation of Serine 2 in the carboxyl terminal domain. RNAPII pauses over the terminator during cleavage and polyadenylation before dissociating. Image created with BioRender.

To study transcription *in vivo*, the most common approach has been chromatin immunoprecipitation (ChIP), in which protein-DNA complexes are stabilized through formaldehyde crosslinking and recovered by immunoprecipitation (Solomon et al., 1988). Coupled with next generation sequencing, ChIP-seq has been widely adopted to explore the genome-wide interactions of RNAPII and co-regulators (Barski et al., 2007; Mikkelsen et al., 2007; Welboren et al., 2009). The occupancy of RNAPII over transcribed regions correlates with nascent transcription. Exonuclease foot printing of RNAPII over DNA (ChIP-exo; Rhee and Pugh, 2012) or RNA (NET-seq; Churchman and Weissman, 2011) and nuclear run-on (PRO-seq; Kwak et al., 2013) have provided high resolution of maps of RNAPII binding to the genome. Together, such methods highlight paused and elongating RNAPII and suggest that very little RNAPII is associated with the promoter in the preinitiation state (Core et al., 2012).

The dynamics of RNAPII transcription *in vivo* has also been explored by tracking single molecules of RNAPII (or co-regulators) or individual transcripts. Such experiments offer a different view of transcription. Fluorescence recovery after photobleaching (FRAP) over arrays of inducible reporter genes reveals that a small fraction (∼13%) of the RNAPII molecules that assemble at the promoter initiates transcription (Darzacq et al., 2007; Stasevich et al., 2014). Monitoring the production of single molecules of mRNA from either such arrays or single genes suggests that RNAPII elongation rate is ∼1000-3000 bp/min and that termination is associated with a prolonged pause (50-70s; Larson et al., 2011; Zenklusen et al., 2008). Single molecule tracking of RNAPII and GTFs reveals that ∼40% of RNAPII is chromatin-associated and that when initiation is blocked, the dwell time of RNAPII (presumably at the promoter) is ∼ 10s (Nguyen et al., 2021). Because these observations would predict that RNAPII levels at the promoter and terminator (as well as pausing sites) should be higher than those over the transcribed region, they are difficult to reconcile with the RNAPII enrichments observed by ChIP-seq.

Single molecule tracking of ssTF and RNAPII binding to enhancers and promoters *in vitro* offers another important perspective. In yeast nuclear extracts, ssTF binding to enhancers (also called upstream activating sequences, UASs) has been observed. Consistent with the consensus model, ssTFs stimulate RNAPII and PIC recruitment to a neighboring promoter (Rosen et al., 2020). Surprisingly, RNAPII and certain PIC components are recruited by ssTFs even in the absence of a promoter (Baek et al., 2021). This suggests that RNAPII is recruited to enhancers/UASs by ssTFs, perhaps through interactions with Mediator, which allows efficient promoter loading of PIC components. However, the association of RNAPII and PIC factors with UASs has not been observed by ChIP-seq.

An alternative to ChIP is chromatin endogenous cleavage (ChEC), in which endogenous proteins of interest are tagged with micrococcal nuclease (MNase; Schmid et al., 2004). Their association with the genome can be monitored by permeabilizing cells and addition of calcium to activate MNase (Schmid et al., 2004). The cleavage events can be identified by next generation sequencing (ChEC-seq2; VanBelzen et al., 2024; Zentner et al., 2015). For ssTFs and nuclear pore proteins, ChEC-seq2 gives results very similar to ChIP-seq or ChIP-exo (Ge et al., 2024; VanBelzen et al., 2024). Likewise, ChEC-seq2 with co-activators and Mediator resembles ChIP (Bruzzone et al., 2018; Grünberg et al., 2016; Saleh et al., 2022). However, we find that ChEC-seq2 with RNAPII gives a pattern of enrichment that was notably different from that observed using ChIP-seq. Whereas ChIP shows strong enrichment of RNAPII over the transcribed region and little enrichment over the promoter or upstream, ChEC-seq2 showed strong enrichment of RNAPII over the promoter, UAS and 3’UTR and little signal over the transcribed region. The ChEC-seq2 enrichment of RNAPII over promoters correlated with both nascent transcription (as measured by SLAM-seq; Herzog et al., 2017) and ChIP-seq enrichment of RNAPII over coding regions, suggesting that it reflects active RNAPII. RNAPII association with UAS regions was strongest for genes that recruit co-activators and was dependent on ssTFs.

The occupancy of RNAPII over UASs and promoters from ChEC-seq2, combined with published RNAPII dynamics, allowed us to develop a Stochastic model for the global kinetics of RNAPII transcription. This model and ChEC-seq2 data offer insight into the effects of genetic perturbations that block transcription globally and suggests that the nuclear pore complex promotes transcription by stabilizing promoter-associated RNAPII. This work suggests that ChEC captures important regulatory events associated with transcription that are missed by ChIP.

## Results

### ChEC-seq2 and ChIP-seq in *S. cerevisiae* yield distinct RNAPII enrichment patterns

To assess ChEC-seq2 with RNAPII, MNase was inserted at the carboxyl terminus of the endogenous genes encoding the RNAPII subunits Rpo21 (also called Rpb1) and Rpb3 (Zentner et al., 2015). These yeast strains, along with a control strain expressing soluble, nuclear MNase (sMNase) were grown in rich medium, harvested and permeabilized to induce MNase activity. Genomic DNA was prepared and converted into ChEC-seq2 libraries (VanBelzen et al., 2024). For comparison, we selected a high-quality RNAPII ChIP-seq dataset from cells grown in rich medium Rpb1 (Vijjamarri et al., 2023b); GEO Accession GSE220578) that used the 8WG16 antibody (Thompson et al., 1989), which recognizes the carboxyl terminal domain of Rpb1 (Philip Komarnitsky et al., 2000). Finally, to confirm that the MNase fusions did not affect RNAPII association with the genome we also generated ChIP-seq data using this antibody from the yeast strains with and without Rpb1-MNase. Over transcriptionally active genes like *ILV5*, ChIP-seq gave strong enrichment of Rpb1 over the transcribed region and terminator and low enrichment over the enhancer/upstream activating sequence (UAS) and the promoter (Figure 2A. 1^st^ row). In contrast, ChEC-seq2 with either Rpb1 or Rpb3 showed strong enrichment at the UAS, promoter, and terminator of *ILV5* and a low enrichment over the transcribed region (Figure 2A, 2^nd^ and 3^rd^ rows; compare with sMNase in black). However, over the repressed *GAL1-10* locus, both ChIP-seq and ChEC-seq2 show background enrichment for RNAPII (Figure 2A, right). Notably, sMNase cleavage over *GAL1-10* reflects both unprotected linkers between well-positioned nucleosomes and nucleosome depletion upstream of promoters (Chereji et al., 2019; Lee et al., 2004); Figure 2A, right). This pattern was unrelated to the trimming of mapped reads to the first base pair (untrimmed tracks in Figure 2 – supplement 1A), the normalization of transcript length used in metagene plots (enrichment over promoters and the 5’ end of genes in Figure 2 - supplement 1B; see Methods), or the presence of the MNase fusion (Figure 2 – supplement 2). Globally, while both ChIP-seq and ChEC-seq2 showed positive Spearman correlation with nascent transcription (measured by SLAM-seq), different regions of genes correlated best with nascent mRNA (Figure 2 – supplement 1C). Nascent transcription correlated best with the enrichment of RNAPII over the promoter from ChEC-seq2 and the enrichment of RNAPII over the transcribed region and terminator from ChIP-seq. Thus, both ChIP-seq and ChEC-seq2 with RNAPII show enrichments that correlate with transcriptional activity, but these two methods reveal complementary interaction patterns.

**Figure 2.**
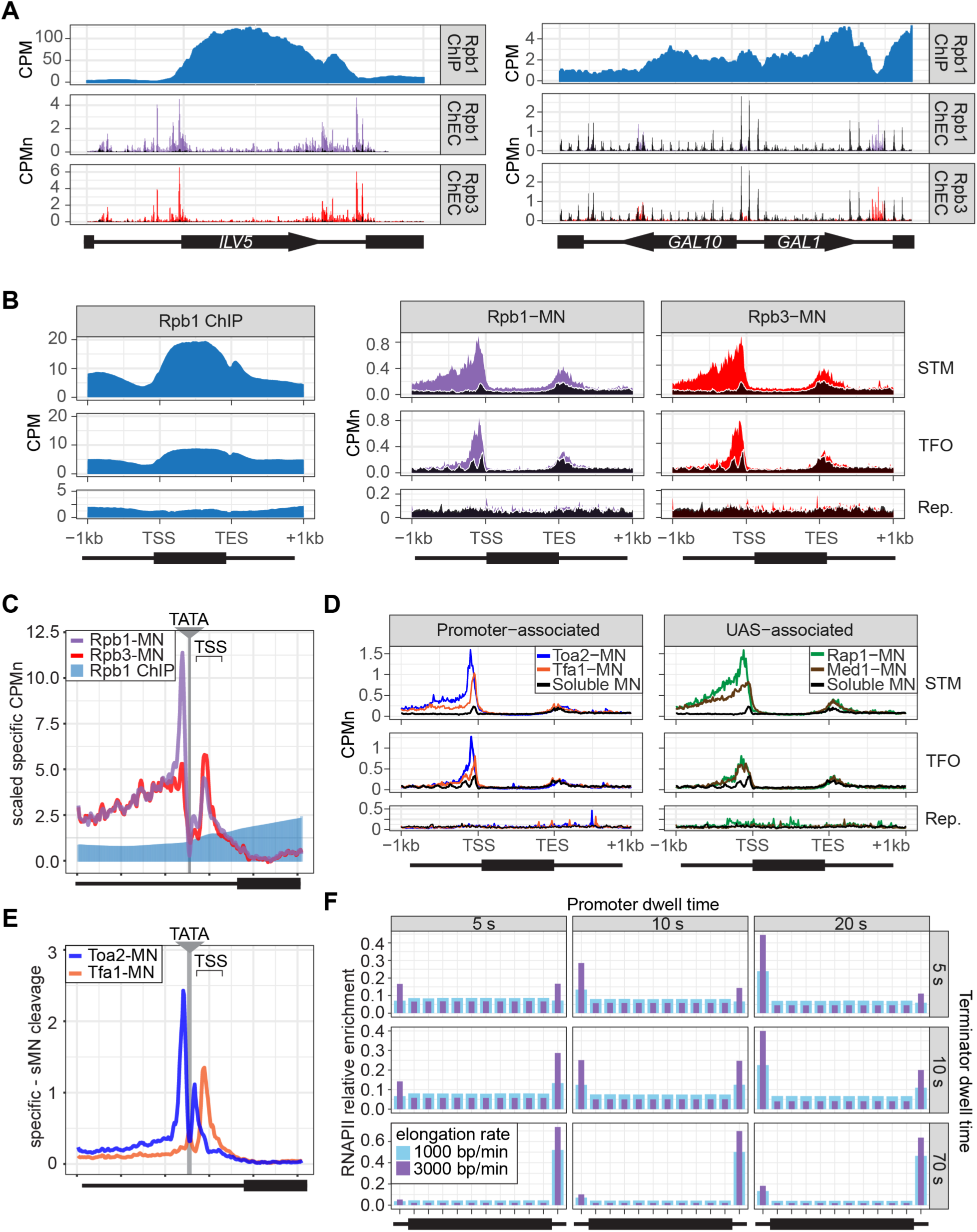
ChEC-seq2 and ChIP-seq reveal distinct RNAPII interactions with the genome. **(A)** Gene plots displaying mean counts per million reads (CPM; Vijjamarri et al., 2023a) for ChIP-seq for Rpb1; or CPM normalized cleavage frequency (CPMn) for ChEC-seq2 for Rpb1-MN or Rpb3-MN over the *ILV5* and *GAL1-10* loci ± 1kb. Plots are smoothed using a window of 10 bp and a step size of 5 bp. Signal from Soluble MNase (sMNase) is shown in black. **(B)** Metagene plots showing average signal over subsets of genes with distinct expression levels and mechanisms of regulation. The average signal from 150 genes with highest expression from STM and TFO classes (Rossi et al., 2021) and 84 repressed genes is plotted, along with sMNase (black; genes listed in Supplementary Table 1). A length-normalized transcript (rectangle), 1 kb upstream of the TSS, and 1 kb downstream of the TES is shown. Rpb1 ChIP-seq (left), ChEC-seq2 with Rpb1-MN (middle) or Rpb3-MN (right). **(C)** ChIP (Rpb1) and ChEC (Rpb1-MN and Rpb3-MN) signal over 597 TATA boxes from expressed genes ± 250bp (Supplementary Table 1; Rhee and Pugh, 2012). The location of the TATA sequence is indicated with a grey bar and the TSS is 50bp ± 39bp to the right of the center of the TATA. For ChEC data, sMNase was subtracted from the respective specific CPMn. **(D)** Metagene plots as in (B) from ChEC-seq2 with Toa2-MN (TFIIA) and Tfa1-MN (TFIIE; left) or the ssTF Rap1-MN and Med1-MN (Mediator; right) from 287 STM genes containing Rap1-peaks, top 150 expressed TFO genes, or 84 repressed genes. **(E)** Mean CPMn at TATA-genes as in (C) from ChEC-seq2 with Toa2-MN (blue) or Tfa1-MN (orange). Panels **A**-**E**, the averages from 3 biological replicates. **(F)** Predicted occupancy of RNAPII based on a range of promoter dwell times (5 - 20 s), elongation rates (1000 - 3000 bp/min) and termination times (5 - 70 s). The transcribed region is 1200 bp divided into 10 x 120bp bins, flanked by an upstream promoter bin and downstream terminator bin. RNAPII occupancy was simulated using a minimal stochastic model. RNAPII was assumed to be immediately present at the promoter and progressed to the transcript region with a rate inverse to the promoter dwell time. It then progressed along a 1200 bp coding region with the indicated elongation rate and terminated transcription with a rate inverse to the terminator dwell time.

Different classes of RNAPII-transcribed yeast genes show distinct mechanisms of transcriptional regulation (Rossi et al., 2021). To more precisely define the differences between ChIP and ChEC, we compared ChIP-seq with ChEC-seq2 over three such classes: 1) genes that bind sequence-specific transcription factors (ssTFs) and coactivators such as <u>S</u>AGA, <u>T</u>up1, <u>M</u>ediator, SWI/SNF (STM), 2) genes bound to ssTFs but not coactivators (transcription factors <u>o</u>nly, TFO) and 3) a set of 330 genes that showed no detectable nascent transcription (repressed). Because these different classes of genes are expressed at different levels (Figure 2 – supplement 1D), we focused on the most highly expressed 150 genes from the STM and TFO classes. Metagene plots of mean RNAPII ChIP-seq over each of these sets of genes reveal strong enrichment over the transcribed region for the STM genes and, to some extent, for the TFO genes, with a notable dip over the promoter (Figure 2B, left). Metagene plots of RNAPII ChEC-seq2 showed a strong enrichment over the promoter for both STM and TFO genes and over the UAS for STM genes (Figure 2B, middle & right). RNAPII was not enriched over repressed genes by either method.

To better understand the ChEC patterns upstream of transcription start sites, mean cleavage by RNAPII was plotted at higher resolution by aligning to 597 high-confidence TATA boxes upstream of expressed genes (based on SLAM-seq), oriented so that the TSS is 50bp ± 39bp to the right (Figure 2C; ± 250bp). Because sMNase cleaves the TATA boxes strongly (Figure 2-supplement 1E) - reflecting either increased accessibility or the T/A sequence preference of sMNase (Dingwall et al., 1981; Horz et al., 1981) - we subtracted the sMNase cleavage from specific cleavage frequency (Figure 2C). Both Rpb1-MN and Rpb3-MN produced cleavage peaks ∼17bp upstream and ∼34bp downstream of the TATA box, although their relative intensities were different (Figure 2D). In contrast, Rpb1 ChIP-seq signal was low over the TATA and TSS (Figure 2C).

The ChEC-seq2 signal for RNAPII over the UAS region correlates with recruitment of coactivators upstream of STM genes, but not upstream of TFO genes (Figure 2B, middle & right), arguing that it is not an artifact of nearby promoters or genes. To better understand the ChEC-seq2 signal over promoters and UAS regions, we mapped proteins expected to interact with the promoter (preinitiation complex (PIC) components TFIIA (Toa2) and TFIIE (Tfa1)) or the UAS (the Rap1 ssTF and Mediator). For this comparison, we selected 287 STM genes near high-confidence Rap1 sites (VanBelzen et al., 2024). While the PIC complex interacted strongly with the promoter region of both STM and TFO genes, Rap1 and Mediator interacted strongly with the UAS region of STM genes (Figure 2D). Rap1 and Mediator also showed a low level of enrichment upstream of the promoter region of TFO genes (Figure 2D). Thus, ChEC-seq2 showed promoter enrichment of PIC components and UAS enrichment of TFs and Mediator.

When mapped over TATA sites, TFIIA (Toa2-MN) produced a major cleavage peak ∼12bp upstream and a minor peak ∼12bp downstream from the TATA box (Figure 2E). TFIIE (Tfa1-MN) showed the strongest peak ∼34bp downstream of the TATA (Figure 2E). These data suggest that ChEC-seq2 reflects the arrangement of TFIIA, RNAPII and TFIIE within the preinitiation complex: TFIIA interacts with DNA immediately upstream of TBP, RNAPII binds on both sides of TBP and TFIIE binds downstream of TBP (see Supplementary movie 1; Aibara et al., 2021; He et al., 2013; Schilbach et al., 2021). Also, consistent with an ordered assembly of the PIC, the peak of TFIIA cleavage 12bp downstream of the TATA box is absent in the RNAPII and TFIIE ChEC data, suggesting that TFIIA binds before RNAPII and TFIIE during PIC assembly and that this site becomes protected when RNAPII and TFIIE join (Supplementary movie 1). Together, these data suggest that ChEC-seq2 captures both UAS-associated RNAPII and the preinitiation complex.

Given the dramatic difference between ChEC-seq2 and ChIP-seq, we next asked if either pattern is consistent with the dynamics of transcription as described in the literature. Because *S. cerevisiae* lacks promoter-proximal pausing (Booth et al., 2016) and has few intron-containing genes that require splicing (Stajich et al., 2007), these slow elongation steps are expected to be absent. Therefore, RNAPII initiation and pausing during termination (Hyman and Moore, 1993) would represent relatively slow steps compared with the rate of elongation. Both *in vivo* and *in vitro* studies in yeast suggest promoter dwell times in the range of approximately 5-20 s (Baek et al., 2021; Nguyen et al., 2021) a termination time of up to 70 s and an elongation rate between 1000-3000 bp/min (Larson et al., 2011; Zenklusen et al., 2008). Using these ranges, we calculated the predicted RNAPII occupancy over the promoter, the transcribed region and the terminator for the typical transcribed yeast gene (see Methods; median size of transcribed region = 1.2kb; Pelechano et al., 2013). Of 24 combinations of dwell times and elongation rates tested, 21 predicted higher occupancy at the promoter than over the transcribed region and 21 predicted higher occupancy at terminators than over the transcribed region (Figure 2F). While some combinations predicted a relatively flat distribution across the gene with lower levels in the promoter, none of the 24 predicted the strong signal over the transcribed region with promoter depletion characteristic of ChIP-seq. Only very short promoter dwell times (*i.e.,* < 1s), produced the low promoter occupancy seen in ChIP-seq (Figure 2 – supplement 1F). This suggests that ChIP-seq is unable to detect functionally important RNAPII interactions at the promoter and UAS that are detected ChEC-seq2.

### ChEC-seq2 detects elongating and phosphorylated RNAPII

Next, we performed ChEC-seq2 with the kinases involved in initiation and elongation, as well as the elongation factor Spt5 (part of DSIF). Phosphorylation of the carboxy terminal domain (CTD) of RNAPII regulates its activity and the association of factors involved in splicing, histone modification, RNA processing. Initiation correlates with phosphorylation of Ser5 of the CTD by Kin28 (Cdk7/TFIIH kinase; Philip Komarnitsky et al., 2000). Elongation is coupled with phosphorylation of Ser2 by Ctk1 (P-TEF-b; CTDK-I; Cdk9; Cho et al., 2001) and Bur1 (P-TEFb; Qiu et al., 2009), and the association of Spt4/5 (DSIF; Hartzog et al., 1998).

Kin28-MN, Ctk1-MN and Spt5-MN showed strong cleavage over active genes and little enrichment over inactive genes (Figure 3A). All three proteins showed maximum cleavage over the promoters of active genes. Kin28 showed significant enrichment over the UAS region of STM genes that was absent from TFO genes (Figure 3A, left). The elongation factor Spt5 showed enrichment over both as well as the transcribed region (Figure 3A, right). In contrast, Ctk1-MN cleavage was primarily localized to promoters (Figure 3A, middle). Higher resolution mapping aligned to TATA boxes confirmed that, while Rpb1 shows peaks of cleavage upstream and downstream of TATA, Kin28, Ctk1 and Spt5 show a single peak downstream, near the TSS (Figure 3B). Furthermore, the signal upstream of the TATA was greatest for Kin28, followed by Ctk1 and then Spt5 (Figure 3B). This suggests that, while Rpb1 shows interactions at the TSS and upstream, factors involved in initiation and elongation are more enriched with the TSS and over the transcribed region.

**Figure 3.**
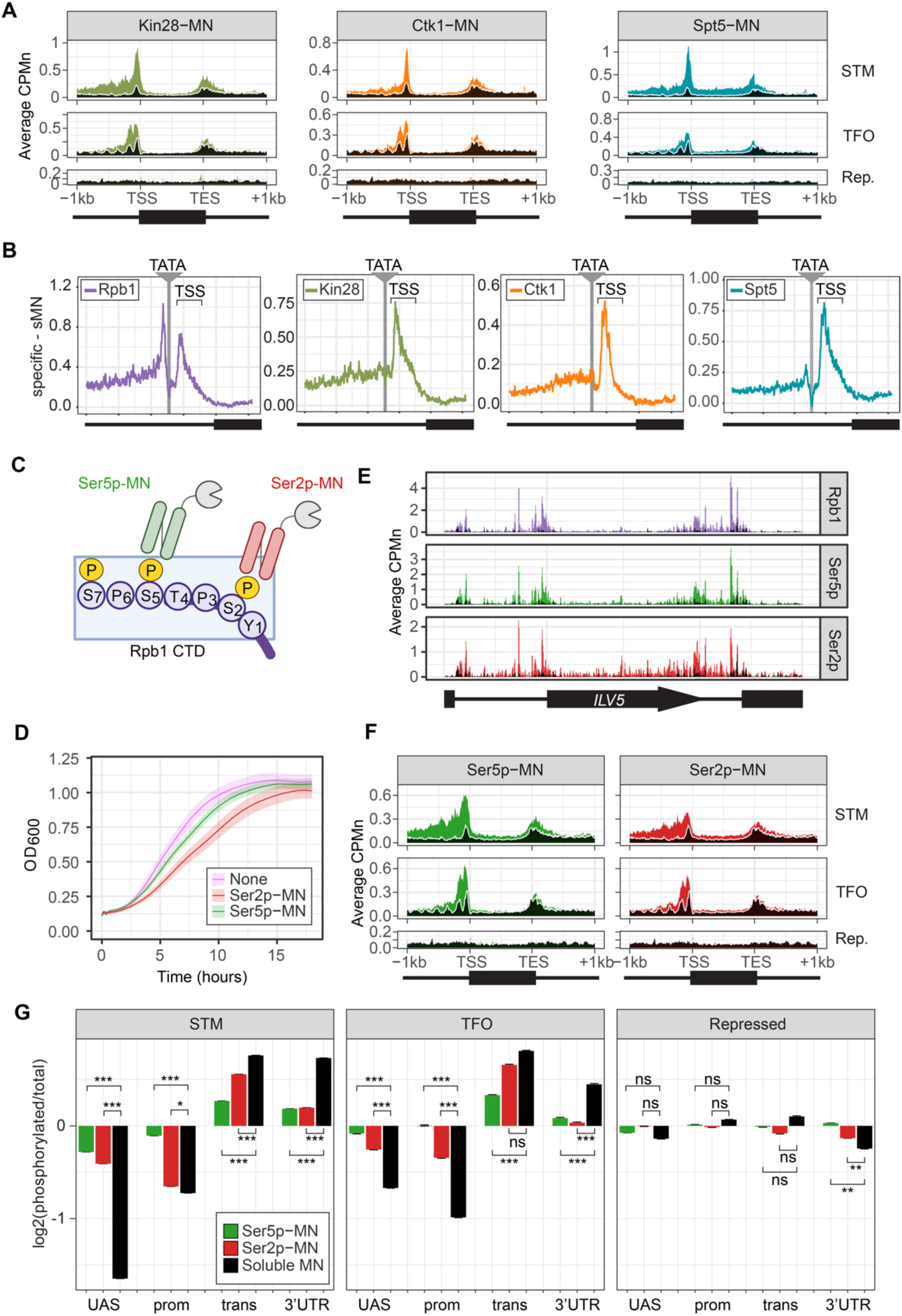
ChEC-seq2 to monitor initiation and elongation. **(A)** Metagene plots showing average CPMn for ChEC with the indicated proteins over subsets of genes with distinct expression levels and mechanisms of regulation. The average signal from 150 genes with highest expression from STM and TFO classes (Rossi et al., 2021) and 84 repressed genes are plotted (Supplementary Table 1). Signal from the sMNase control is shown in black with a white line for contrast. **(B)** Average CPMn for the indicated proteins over 597 TATA boxes ± 250 bp from expressed genes (Supplementary Table 1; Rhee and Pugh, 2012). The location of the TATA sequence is indicated with a grey bar and the TSS is 50bp ± 39bp to the right of the TATA. The signal for sMNase was subtracted from each. **(C)** Schematic for mintbody-MNase constructs. Two single chain variable fragments of IgG specific to phosphorylation of Serine 5 (Ser5p) or Serine 2 (Ser2p) of the CTD of RNAPII (Ohishi et al., 2022; Uchino et al., 2021) were tagged at their C-termini with MNase. Created with BioRender. **(D)** OD_600_ of the parent strain (pink), strain expressing Ser5p-MN (green), and strain expressing Ser2p-MN (red) over time in culture (average ± standard deviation). **(E)** Average CPMn from ChEC-seq2 with Rpb1-MN (purple), α-Ser5P-MN (green) and α-Ser2P-MN (red) at *ILV5* ± 1kb. Plots were smoothed with a step size of 5 and window of 10. **(F)** Metagene plots as in (A), but with signal from Ser5p-MN (green; left) and Ser2p-MN (red; right). **(G)** For each protein, the relative enrichment at UAS, promoter, transcript, and 3’UTR regions was calculated and normalized by region length for each gene. The resulting values were normalized to those values for Rpb1-MN and the average from all genes in each group is plotted. Error bars represent the estimated variance between biological replicates from standard deviation (n = 3). Differences between ratios and estimated variance were used to calculate a z score and p-value; *p<0.05, **p<0.01, ***p<0.001. All panels represent the average from 3 biological replicates.

To confirm that the ChEC cleavage pattern by Kin28 and Ctk1 reflects their activity, we developed a method to measure RNAPII phosphorylation by ChEC-seq2. Two single chain IgG fragments that recognize phosphorylated Ser2 (Ser2p) RNAPII CTD or phosphorylated Ser5 (Ser5p) RNAPII CTD (Mintbodies) have been expressed as GFP- and SNAP-tagged fusions and shown to localize at transcriptionally active loci in mammalian cells (Ohishi et al., 2022; Uchino et al., 2021). We constructed Mintbody-MNase (Mb-MN) fusions to detect these phosphorylated forms of RNAPII (Figure 3C; α-Ser2p-MN and α-Ser5p-MN). Because binding phosphorylated CTD could compete for critical interactions with RNAPII, we tested several promoters to identify an expression level that produced the smallest growth defect (not shown). Strains expressing the Mb-MNs from the *ADH1* promoter had a minimal growth defect (Figure 3D) and cleaved chromatin upon permeabilizing cells and addition of calcium (Figure 3-supplementary Figure S2A). Both α-Ser5p-MN and α-Ser2p-MN give patterns very similar to those produced by their respective kinases; Ser5p was more enriched over promoters and UAS regions, while Ser2p was more evident in the transcribed region (Figure 3E & F). To better compare these patterns, we normalized mean cleavage over promoters, UAS regions, transcribed regions and 3’UTR regions by each Mb-MN (or sMNase) to that by Rpb1 (Figure 3G). Ser5p and Ser2p levels were lower than Rpb1 over the UAS and promoter, but higher than Rpb1 over the transcript and 3’UTR (Figure 3G). Furthermore, consistent with their patterns observed by ChIP, the levels of Ser2p were lower than those of Ser5p over the UAS and promoter and higher than those of Ser5p over the transcript (Figure 3G). Thus, ChEC-seq2 can reveal RNAPII recruitment, initiation and elongation during transcription.

### Global transcriptional changes are detected by ChEC-seq2

To further validate the biological significance of RNAPII ChEC-seq2, we examined the effects of an environmental perturbation that results in a large-scale transcriptional change. Cells exposed to 10% ethanol in growth medium show widespread changes in transcription, downregulating hundreds of genes enriched for those involved in ribosome biogenesis (GO: 0042254; blue in Figure 4A) and upregulating genes enriched for chaperones (GO: 0009266; red in Figure 4A). ChEC-seq2 using Rpb1-MN, Kin28-MN, Ctk1-MN, α-Ser5p-MN and α-Ser2p-MN captures these changes. These proteins showed increased enrichment over the *HSP104* chaperone gene following ethanol treatment (Figure 4B). Likewise, metagene plots over the top 100 induced genes showed increased cleavage by Rpb1-MN, Kin28-MN, Ctk1-MN as well as their products Ser5p and Ser2p upon ethanol treatment (Figure 4C & D, left). Notably, over the transcribed region, enrichment was higher at the 3’ end, especially for Ser2p and Ctk1-MN (Figure 4C & D, left). In contrast, metagene plots of the average change in cleavage over the 137 ribosomal protein genes showed strong decreases in cleavage by all of these proteins (Figure 4C & D, right). The changes in sMNase cleavage were generally the opposite of what we observed with the specific proteins (Figure 4C & D, black trace/column). Thus, ChEC-seq2 can capture biologically relevant changes in RNAPII association, its regulators and its phosphorylation states that reflect large-scale changes in global transcription.

**Figure 4.**
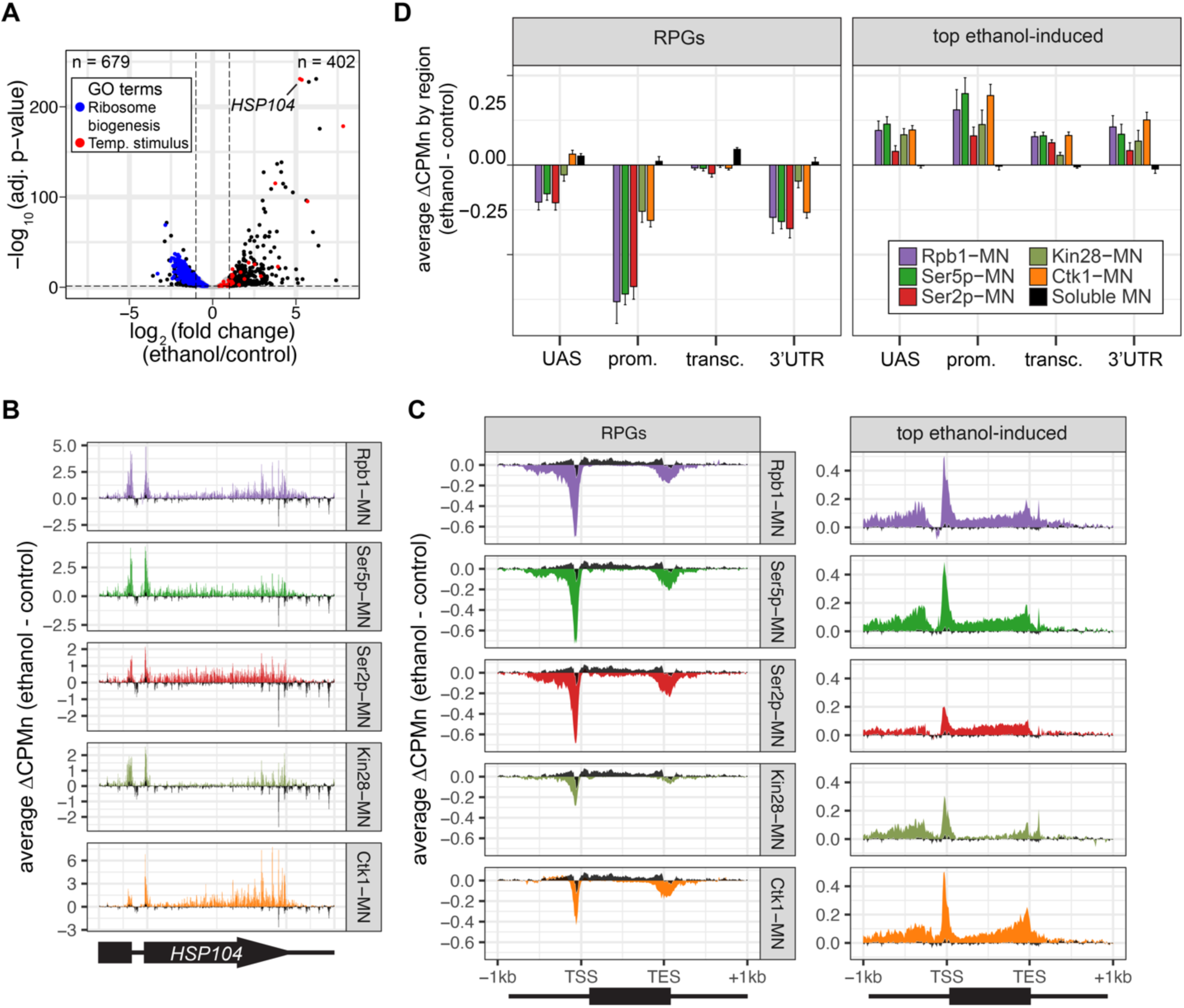
Transcriptional response to ethanol stress results in widespread changes in RNA polymerase II ChEC-seq2. **(A)** Volcano plot of fold change vs. −log_10_ of adjusted p-values of 5295 mRNAs comparing cells treated with 10% ethanol for 1 h vs. untreated cells. The mRNAs belonging to the most statistically significant terms from Gene Ontology Enrichment analysis of the 402 upregulated mRNA (response to temperature stimulus, GO:0009266; red) or 679 downregulated mRNA (ribosome biogenesis, GO:0042254; blue) are highlighted. **(B)** Average change in CPMn over *HSP104* ± 1kb (ethanol – untreated) from ChEC-seq2 with Rpb1-MN, mintbody-MNase constructs (α-Ser5p -MN, α-Ser2p-MN), Kin28-MN, and Ctk1-MN. Data were smoothed with a step size of 5 and window of 10. **(C)** Metagene plots showing the average change in CPMn (ethanol – untreated) from ChEC-seq2 of the top ethanol-induced genes (100 genes, right; Supplementary Table 1) and the downregulated ribosomal protein genes (137 genes, left; Supplementary Table 1). CPMn for sMNase is shown in black. **(D)** Gene-region enrichment of each protein relative to Rpb1-MN. For each protein and each gene, the average change in CPMn (ethanol – untreated) was calculated. Signal was binned into gene regions and normalized by region length. The length-normalized region signal relative to total signal (all gene regions) was calculated. The average for 137 RPGs and the top 100 ethanol-induced ± SEM is plotted. For all panels, the average from 3 biological replicates is shown.

### RNAPII ChEC-seq2 upon chemical-genetic perturbations of transcription

Next, we tested the effect of blocking either PIC formation or initiation on RNAPII/PIC occupancy by ChEC-seq2. PIC formation was blocked by depleting TFIIB using auxin-induced degradation (Sua7-AID; Figure 5A) and initiation was inhibited by treating an analog-sensitive allele of Kin28 with the ATP analog CMK (*kin28-is*; Rodríguez-Molina et al., 2016). These treatments resulted in strong down-regulation of nascent transcription (SLAM-seq; Figure 5B) and inhibition of growth (Figure 5 - supplement 1), respectively. ChEC-seq2 with Rpb1 following 20 minutes of depletion of TFIIB showed a clear decrease of Rpb1-MN cleavage over the promoters of the 150 most highly transcribed STM and TFO genes (Figure 5C). TFIIB depletion caused a shift in sMNase cleavage from the TSS downstream (Figure 5C). Neither Rpb1-MN nor sMNase cleavage over repressed genes was altered by TFIIB depletion (Figure 5C). This suggested that Rpb1 occupancy over the promoters of STM and TFO genes requires TFIIB.

**Figure 5.**
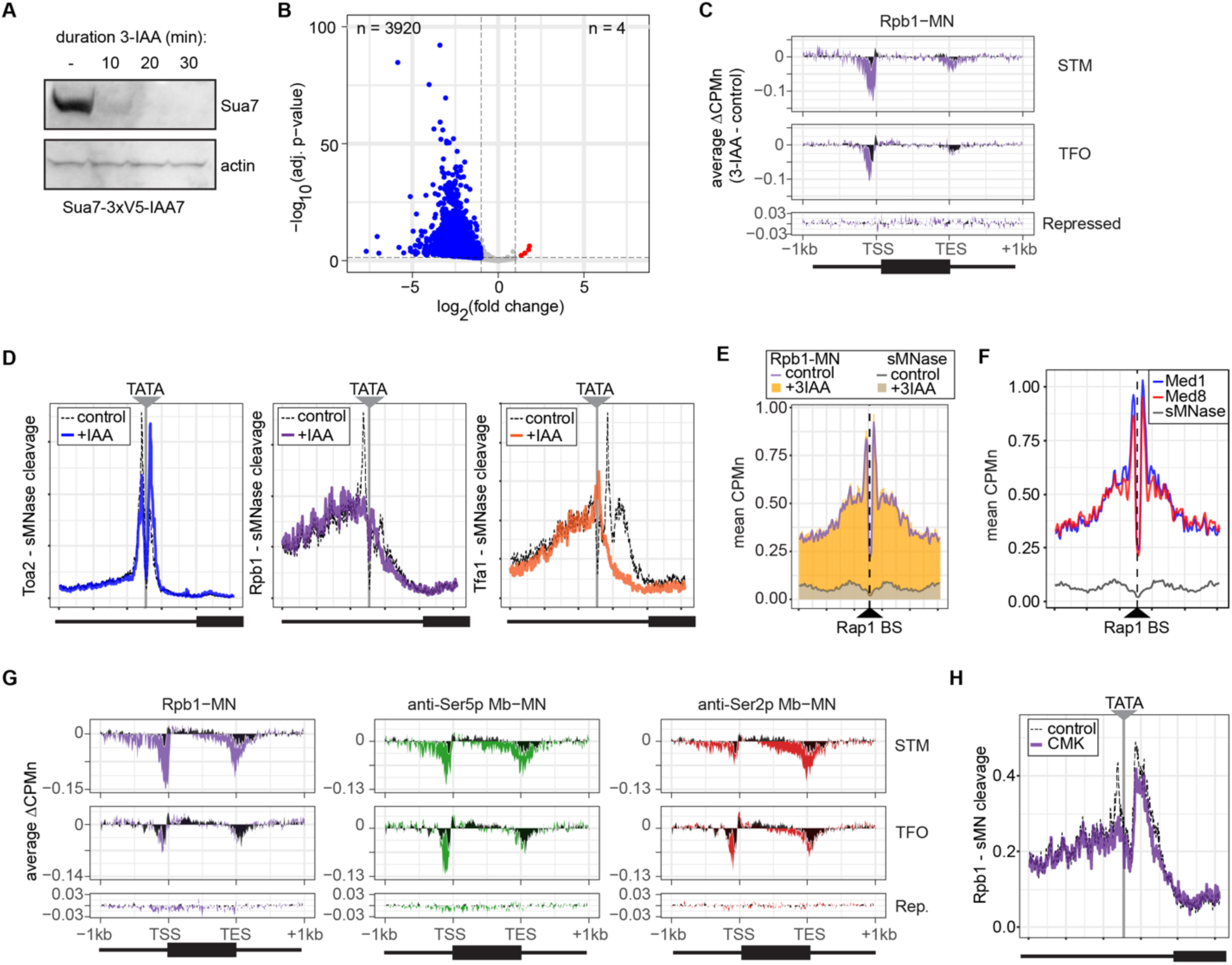
Conditional depletion of TFIIB and inhibition of TFIIH kinase cause distinct eRects on promoter-associated RNAPII. **(A)** Chemiluminescent immunoblot of Sua7-3V5-IAA7 at the indicated time points following addition of 3-IAA. Signal from actin is shown as a loading control. **(B)** Volcano plot of log_2_ fold change in nascent RNA vs the −log_10_ of the adjusted p-value following degradation of Sua7-3V5-IAA7 via treatment with 3-IAA for 60 minutes. Of 5295 mRNAs, 3920 mRNAs were significantly decreased (blue; LFC ≤ −1 & adj. p < 0.05) and 4 mRNAs were significantly increased (red; LFC ≥ 1 & adj. p < 0.05). Cells were grown in synthetic complete medium. **(C)** Metagene plots showing the average change in CPMn (3-IAA – control) from ChEC-seq2 with Rpb1-MN upon degradation of Sua7-3V5-IAA7 for 20 minutes over 150 genes with highest expression from STM and TFO classes (Rossi et al., 2021) and 84 repressed genes are plotted (Supplementary Table 1). Cells were grown in SDC. **(D)** Metasite plot over the TATA boxes ± 250bp from 597 expressed, mRNA-encoding genes (Supplementary Table 1; Rhee and Pugh, 2012). In each case, sMNase CPMn was subtracted from the specific CPMn and the untreated control is shown in grey for comparison. The location of the TATA sequence is indicated with a grey bar. Cells were grown in YPD and Sua7-3V5-IAA7 was depleted for 60 min. **(E)** Metasite plot of average CPMn from Rpb1-MN and sMNase cleavage over 896 high-confidence Rap1 sites (VanBelzen et al., 2024). Purple and dark grey lines represent mean Rpb1-MN and sMNase cleavage in untreated cells; orange and grey columns represent mean Rpb1-MN and sMNase cleavage upon Sua7 depletion for 60 min. **(F)** Metasite plots of average CPMn from Med1-MN (brown), Med8-MN (magenta), sMNase (grey) over Rap1 sites as in (E). **(G)** Metagene plots as in (**C**) of average change in signal (CMK – control) upon inhibition of *kin28is* for Rpb1-MN (purple), Ser5p-MN (green), Ser2p-MN (red). Cells were grown in SDC and treated with 5 µM CMK for 60 minutes. **(H)** Metasite plot over TATA boxes as in (**D**) for the sMNase-corrected signal from Rpb1-MN before (grey) and after (purple) inhibition of *kin28-is* with 5 µM CMK for 60 min. For all panels, the average of three biological replicates is plotted.

Higher resolution mapping of RNAPII (Rpb1-MN), TFIIA (Toa2-MN) and TFIIE (Tfa1-MN) cleavage over TATA boxes revealed that, upon TFIIB depletion, TFIIA occupancy shifted from the major upstream peak to the downstream peak (Figure 5D). RNAPII and TFIIE peaks near the TATA and TSS were lost (Figure 5D). This supports the notion that the downstream peak of TFIIA is blocked by RNAPII/PIC binding. Furthermore, the cleavage by Rpb1 upstream of the TATA box was unaffected by depletion of TFIIB (Figure 5D, middle), suggesting that TFIIB is required for proper PIC formation over the promoter, but is not required for association with upstream UAS elements.

To test this hypothesis, we mapped RNAPII (Rpb1-MN) cleavage over 896 high-confidence sites for the ssTF Rap1 (VanBelzen et al., 2024). Rap1 regulates hundreds of highly expressed genes and RNAPII ChEC-seq2 showed strong enrichment flanking Rap1 sites, while sMNase did not (Figure 5E). This correlates with Mediator occupancy (Figure 5F). Depletion of TFIIB had no significant effect on RNAPII occupancy over Rap1 sites (Figure 5E). Thus, RNAPII recruitment to the promoter is dependent on TFIIB, while RNAPII recruitment to the UAS is not.

Inhibition of *kin28-is* with CMK also lead to a strong decrease of RNAPII over the promoter, transcribed region and 3’UTR, especially for the STM genes (Figure 5G). As expected, this was associated with a strong decrease of Ser5 phosphorylation and Ser2 phosphorylation (Figure 5G). Cleavage by α-Ser5p-MN was most strongly decreased at the promoter, while cleavage by α-Ser2p-MN was most strongly decreased at the 3’ end of the transcribed region. No changes in cleavage were observed at repressed genes. RNAPII cleavage over 597 TATA boxes near expressed genes was also decreased upon Kin28 inhibition, but this effect was not as strong as that observed upon depletion of TFIIB (Figure 5H). Thus, inhibition of Kin28 led to an apparent decrease in total RNAPII and its Ser2 and Ser5 phosphorylated forms from highly expressed genes.

### Developing a kinetic model for transcription based on ChEC-seq2 RNAPII occupancy

Because ChEC-seq2 provides information about important regulatory steps that have not been evident from previous global studies, we used these data (as well as ChIP-seq data) to develop models for the global kinetics of yeast RNAPII transcription. Steady-state occupancy of RNAPII should reflect the rates of several steps: RNAPII recruitment to the UAS and/or promoter, PIC assembly, initiation, elongation and termination. We developed a stochastic computational model for these steps (Figure 6A) by fixing rates that have been experimentally determined (*k*_1_, *k*_-1_, *k*_3_, *k*_5_, *k*_6_, *k*_7_; Table 1) and optimizing the remaining rates to fit to the RNAPII occupancy observed from either ChIP-seq or ChEC-seq2. To capture the distinct mechanisms of RNAPII recruitment, we modeled the STM and TFO gene classes separately: for the STM class, we assumed that all RNAPII is recruited first to the UAS (reflecting *k*_1_) before being transferred to the promoter (reflecting *k*_2_); for the TFO class, RNAPII is recruited directly to the promoter (reflecting *k*_3_). In genes with a UAS (*i.e.*, STM genes), RNAPII is recruited nearly exclusively to the UAS through ssTFs and coactivators (Baek et al., 2021), and we therefore omitted RNAPII recruitment to the promoter (*k*_3_) the STM model. We also modeled dissociation from the UAS (reflecting *k*_-1_, STM class) and promoter (reflecting *k*_-3_, both classes), as well as the possibility of reversal from promoter to UAS (reflecting *k*_-2_, STM class).

**Figure 6.**
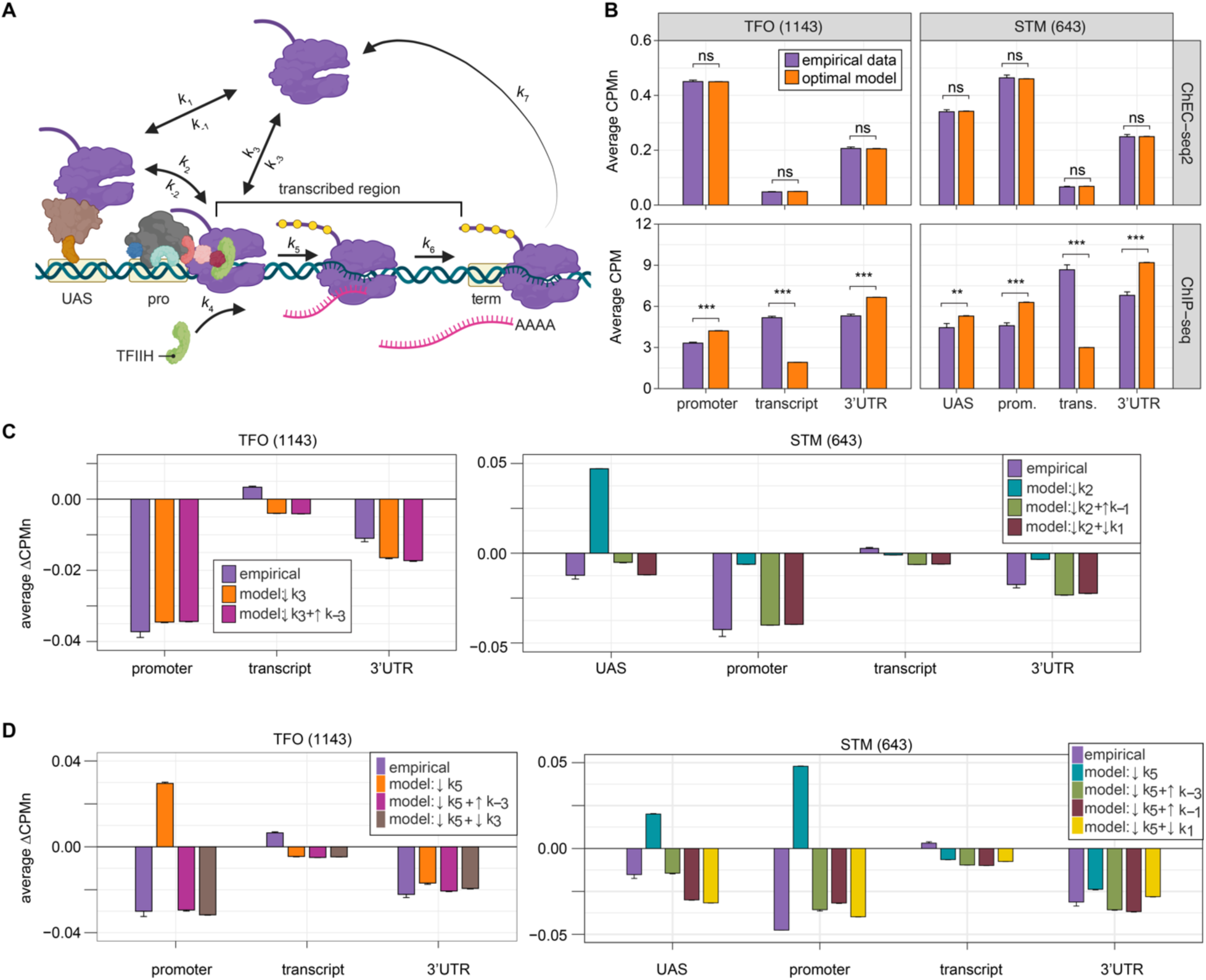
Global kinetic model for RNAPII transcription. **(A)** Schematic for a model of the global kinetics of transcription by RNA Polymerase II in *S. cerevisiae*. Two alternative mechanisms of RNAPII recruitment are shown: 1) direct recruitment to TFO promoters governed by rates *k*_3_ and k_-3_ and 2) recruitment to the STM UASs facilitated by a ssTFs and coactivators such as Mediator (*k*_1_ and *k*_-1_), followed by transfer to the promoters (*k*_2_ and k_-2_). After RNAPII arrives at the promoter it can dissociate at rate *k*_-3_ until TFIIH is recruited (*k*_4_), followed by initiation (*k*_5_). RNAPII elongation (k_6_) across the transcribed region produces mRNA. Pausing during termination is determined by the dissociation rate *k*_7_. Transcription is modeled as a stochastic, processive process with successful recruitment of TFIIH representing the committed step. Rates *k*_4_, *k*_5_, *k*_6_, and *k*_7_ are irreversible. Image created with BioRender. **(B)** The average Rpb1 signal (purple) from ChEC-seq2 (top) and ChIP-seq (bottom) over the indicated regions from 1143 TFO-class genes and 643 STM-class genes that are expressed in SDC (Supplementary Table 1). Rates *k*_2_, *k*_-2_, *k*_-3_, and *k*_4_ were explored to fit the empirical data for each dataset. The remaining rates were drawn from published values (see Table 1). RNAPII occupancy was simulated across gene regions. UAS, Promoter, and 3’UTR were represented by a single 120 bp bin and the transcript region was composed of 10 sequential bins to represent a 1200 bp transcript. The average predicted occupancy for RNAPII over each region from the models (*i.e.* sets of rates) that best matched the empirical data are shown (see Methods). For Rpb1 ChEC-seq2, 789 STM models and 371 TFO models fit the empirical data. Using the same fit-thresholds for ChIP-seq data produced no models. Instead, the average predicted occupancy from an equal number of the top-performing ChIP-seq models as used in ChEC-seq2 simulations (*i.e.*, 789 for STM and 371 for TFO) was used to generate the predictions shown (Table 1). **(C, D)** The average change in Rpb1-MN by ChEC-seq2 at each gene-region following conditional depletion or inactivation of PIC components (purple) for 1143 TFO-class genes and 643 STM-class genes that are expressed in SDC is shown. The rates from the ensemble of best models in (**B**) were adjusted to model the observed changes in Rpb1-MN over each gene region. **(C)** The average change in Rpb1-MN (3-IAA - control) following conditional depletion of TFIIB (purple, from Figure 5C). For TFO-class genes, a decrease in promoter recruitment (τ*k*_3_) with or without an increase in promoter dissociation (τ*k*_3_ + τ*k*_-3_) fit the empirical findings. For STM genes, a decrease in transfer from UAS (τ*k*_2_) combined with an increase in dissociation from UAS (τ*k*_2_ + τ*k*_-1_) or decrease in UAS recruitment (τ*k*_2_ + τ*k*_1_) fit the observed changes (Table 2). **(D)** The average change in Rpb1-MN (CMK - control) following inhibition of TFIIH kinase (purple, from Figure 5G). For TFO-class genes, a decrease in initiation (τ*k*_5_) combined with an increase in dissociation from promoter (τ*k*_5_ + τ*k*_-3_) or decrease in promoter recruitment (τ*k*_5_ + τ*k*_3_) fit the observed changes. For STM, a decrease in initiation (τ*k*_5_) combined with either an increase in dissociation from promoter (τ*k*_5_ + τ*k*_-3_), an increase in dissociation from UAS (τ*k*_5_ + τ*k*_-1_), or a decrease in UAS recruitment (τ*k*_5_ + τ*k*_1_) fit the observed changes (Table 2).

**Table 1.** Model parameters.

| Rate | Published Values | Standard Growth - ChEC-seq |  | Standard Growth - ChIP-seq |  | GCN4 |  | Rate Description |
| --- | --- | --- | --- | --- | --- | --- | --- | --- |
|  |  | Selected Rate | Functional Range | Selected Rate | Functional Range | Selected Rate | Functional Range |  |
| $k_{1, \text{UAS}}$ | 0.0019 - 0.0027/s (Rosen 2020) | 0.002/s | - | 0.002/s | - | 0.002/s | - | Recruitment of RNAPII to UAS |
| $k_{-1, \text{UAS}}$ | 0.001 - 0.005/s (Rosen 2020) | 0.003/s | - | 0.003/s | - | 0.003/s | - | Dissociation of RNAPII from UAS |
| $k_{2, \text{UAS}}$ | None Identified | - | 0.03 - 0.2/s | - | 0.06 - 0.2/s | - | 0.03 - 0.2/s | Transfer of RNAPII from UAS to Promoter. Requirement for assembly early PIC components at Promoter is incorporated |
| $k_{-2, \text{UAS}}$ | None Identified | - | 0.0 - 0.15/s | - | 0.0 - 0.18/s | - | 0.0 - 0.04/s | Transfer of RNAPII from Promoter to UAS |
| $k_{3, \text{Promoter}}$ | None identified. Used $k_1$ | 0.002/s | - | 0.002/s | - | NA | - | Recruitment of RNAPII to Promoter |
| $k_{-3}$ | None Identified | - | 0.0 - 0.03/s | - | 0.0 - 0.03/s | - | 0.0 - 0.03/s | Dissociation of RNAPII from Promoter |
| $k_4$ | None Identified | - | 0.0075 - 0.09/s | - | 0.075 - 0.2/s | - | 0.01-0.15/s | Recruitment of TFIH |
| $k_5$ | 0.1/s (TFIIH residency, Wu 2020) | 0.1/s | - | 0.1/s | - | 0.1/s | - | Initiation. Phosphorylation of RNAPI CTD by TFIH Kinase (Kin28) |
| $k_6$ | 1 - 3 kb/min | 1 kb/min | - | 1 kb/min | - | 1 kb/min | - | Elongation. |
| $k_7$ | 0.009 - 0.034 /s (Larson 2011 on MDN1) | 0.037/s | - | 0.037/s | - | 0.0614/s | - | Termination. |

**Table 2.**
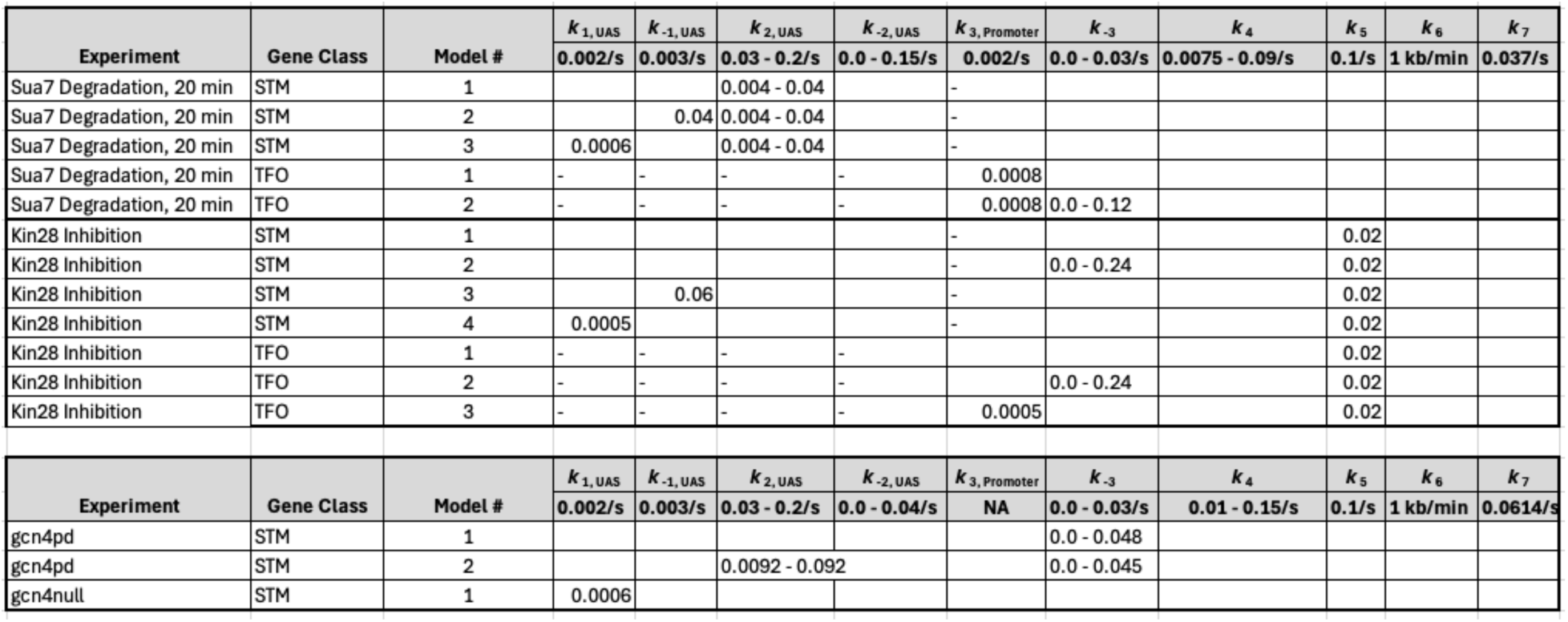
Perturbations of the model.

Fitting to the RNAPII occupancy from ChIP-seq or ChEC-seq over different regions (UAS, promoter, transcribed region or 3’UTR), we identified the optimal range of values for the undefined rates (*i.e.*, *k*_2_, *k*_-2_, *k*_-3_, and *k*_4_), producing an ensemble of best-fit models (Figure S5). Agreement between the models and the data was measured using cosine similarity (Methods). The models trained on the ChEC-seq2 occupancy for either the TFO or STM genes showed excellent agreement with the data (cosine similarity > 0.995; Figure 6B, top & Figure 6 – supplement 1A & C). Optimal agreement between the models and ChEC-seq data was achieved by using the lower bound for dwell time at the terminator from Zenklusen et al., 2008 and Larson et al., 2011 (30 seconds; *k*_7_ = 0.0325 s^−1^; Table 1). Importantly, the rates that are shared between the STM and TFO models are identical (Table 1). Thus, modeling RNAPII occupancy data from ChEC-seq2 produced a range of plausible values for the rates of transcription that agrees well with the empirical data (Table 1).

Using the published rates, neither model was able to find rates for the other steps that produced occupancy that matched that observed by ChIP-seq (*i.e.*, there were no models with cosine similarity > 0.9; Figure 6 – supplement 1B & D). The best ChIP-seq models predicted RNAPII occupancy over all regions that was significantly different from that observed (Figure 6B, bottom). By varying the published rates, the model could produce the occupancies observed by ChIP-seq (Figure 6 – supplement 1E & F). However, this required eliminating dissociation from the promoter (*k*_-3_), increasing the initiation rate (*k*_5_) two-fold with instantaneous recruitment of TFIIH (*k*_4_) and increasing the termination rate ∼4.3-fold above the maximum published rate (*k*_7_ = 0.14 s^−1^; Figure 6 – supplement 1 E, inset table). Thus, although it is possible to model the RNAPII occupancy observed by ChIP-seq, the predicted rates are difficult to reconcile with the literature.

We explored which rates in the model could account for the effects of TFIIB depletion (Figure 6C) and Kin28 inhibition (Figure 6D; Methods) on mean RNAPII occupancy over UASs, promoters, transcribed regions and 3’UTRs. Consistent with a role for TFIIB in recruiting RNAPII to the promoter, reducing the rate of RNAPII recruitment (*k*_3_) to the promoters of TFO genes produced RNAPII occupancy changes that matched the observed effects of TFIIB depletion (Figure 6C, left; Table 1).

For the STM genes, decreasing k_2_ alone (*i.e.*, the rate of RNAPII transfer from the UAS to promoter) predicted an accumulation of RNAPII at the UAS and did not agree well with the data (Figure 6C, right). Instead, models that decreased k_2_ and *either* increased the rate of dissociation from the UAS (*k*_-1_) or decreased the rate of RNAPII recruitment to the UAS (*k*_1_) produced RNAPII occupancies that agreed well with the data (Figure 6C, right; Table 1). Therefore, for STM genes, the model predicts that depletion of TFIIB may both reduce RNAPII recruitment to the promoter and reduce recruitment of RNAPII to, or stimulate RNAPII dissociation from, the UAS.

Next, we asked which rates in our kinetic model could account for the effects of inhibiting Kin28. Modeling a decrease in the rate of initiation (*k*_5_) predicted an accumulation at the promoter (and UASs of STM class genes), which is not observed (Figure 6D). Instead, the effects of inhibiting Kin28 fit best with destabilizing RNAPII bound to the UAS or promoter, either by decreasing recruitment (*k*_1_ or *k*_3_, respectively) or by increasing dissociation (*k*_-1_ or *k*_-3_, respectively; Table 1). Indeed, for TFO class genes, either an increase in promoter dissociation (*k*_-3_) or a decrease in promoter recruitment (*k*_3_) with a decrease in initiation (*k*_5_) produced occupancies that agreed with the data (Figure 6D, left; Table 1). Similarly, for STM class genes, incorporating an increase in promoter dissociation *(k*_-3_), an increase in UAS dissociation (*k*_-1_) or decrease in UAS recruitment (*k*_1_) with a decrease in initiation (*k*_5_) resulted in fits that agreed with the empirical findings (Figure 6D, right). Notably, for STM class genes, the combination of a decrease in initiation with an increase in promoter dissociation produced the best fit at the UAS. This suggests a feedback mechanism between PIC formation and initiation. Together, these findings indicate that the changes in RNAPII occupancy observed by ChEC-seq2 upon perturbation of PIC components can be explained by reasonable changes in transcriptional rates.

### ChEC-seq2 suggests a role for the NPC in stabilizing promoter association of RNAPII

Hundreds of active yeast genes physically associate with the NPC and this is dependent on ssTFs (Ahmed et al., 2010; Brickner et al., 2019, 2012; Casolari et al., 2005, 2004; Light et al., 2010; Randise-Hinchliff et al., 2016; Vosse et al., 2013). Mutations that disrupt this interaction cause a quantitative decrease in transcription (Ahmed et al., 2010; Brickner et al., 2016). For example, a mutation in the Gcn4 TF that blocks interaction with the NPC results in a quantitative decrease in transcription of Gcn4 targets (genes involved in amino acid biosynthesis; Brickner et al., 2019; Hinnebusch and Fink, 1983). This mutation replaces three amino acids within a 27 amino acid Positioning Domain (PD_GCN4_) that does not overlap the activation or DNA binding domains (Brickner et al., 2019). We confirmed this effect by measuring nascent transcription upon amino acid starvation in *gcn4-pd* strains or a wildtype control (Materials & Methods). Although both *GCN4* and *gcn4-pd* mutant strains showed widespread transcriptional changes upon amino acid starvation (Figure 7A), the upregulation (and downregulation) of transcription was quantitatively stronger for the *GCN4* strain (Figure 7A, right panel). We tested if this transcriptional defect is associated with a competitive fitness defect by competing *GCN4* and *gcn4-pd* strains in the absence of histidine ± 3-amino triazole (3-AT, an inhibitor of the His3 enzyme, which selects for maximal expression of *HIS3*). The relative abundance of *GCN4* and *gcn4-pd* strains was quantified using Sanger sequencing (Sump et al., 2022). The *GCN4* strain showed greater fitness under both conditions, but this was particularly evident in the presence of 3-AT (Figure 7B).

**Figure 7.**
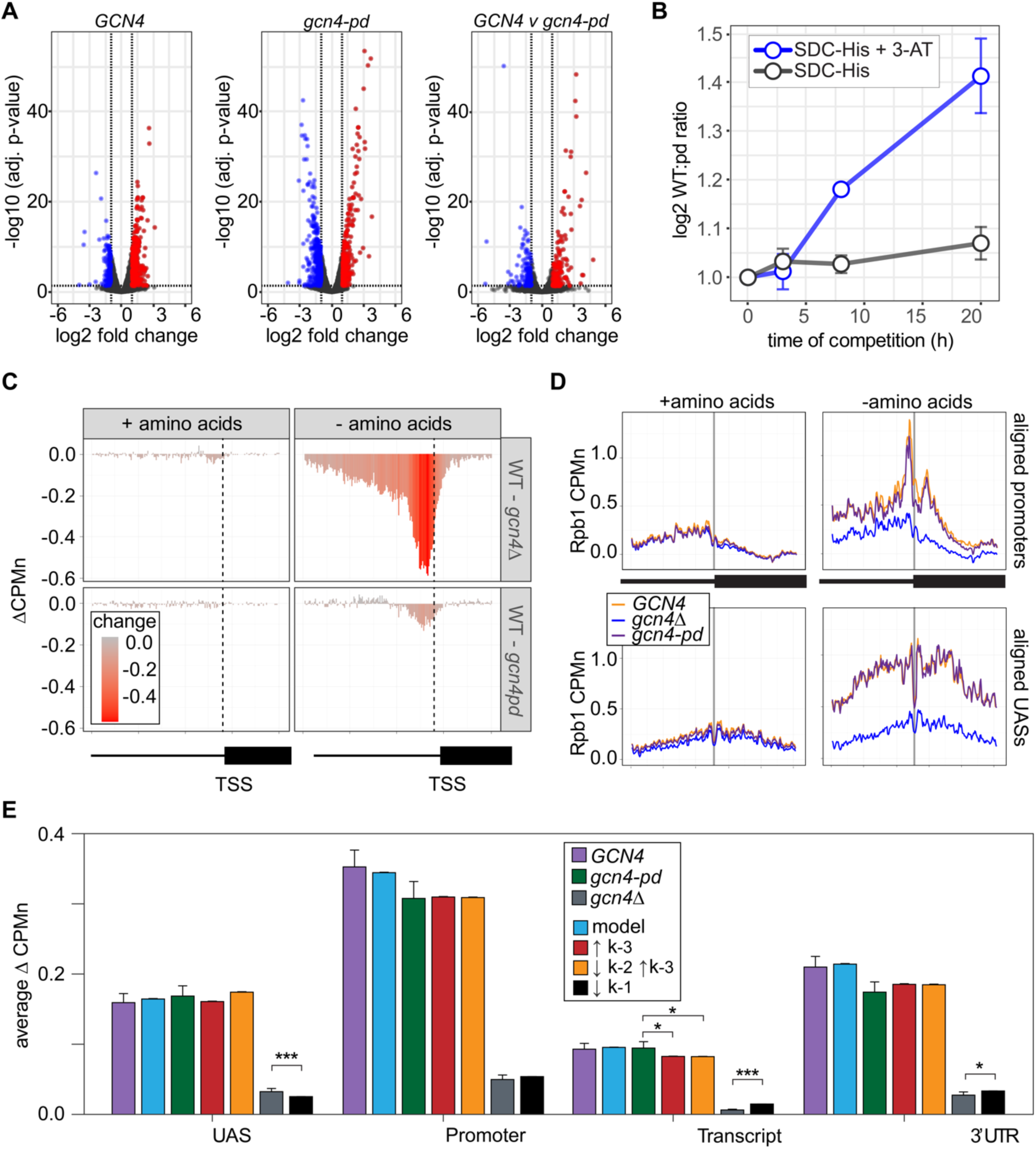
The Gcn4 positioning domain stabilizes RNAPII association with the promoter without aRecting recruitment to the UAS. **(A)** Volcano plot of log_2_ fold-change in nascent RNA vs - log_10_ adjusted p-values from cells starved for histidine for 1h vs. cells in complete medium. Nascent RNA counts for a total of 5295 mRNAs were measured in *GCN4* (left) and *gcn4-pd* (middle). The log_2_ fold-change in nascent RNA vs −log_10_ adjusted p-values in *GCN4* vs. *gcn4-pd* from cells grown in the absence of histidine (right). Significantly downregulated (blue; LFC ≤ −1 & adj. p < 0.05) upregulated (red; LFC ≥ 1 & adj. p < 0.05) genes are highlighted. (**B**) Relative abundance of *GCN4* and *gcn4-pd* strains in a mixed culture, determined by quantifying the relative abundance of the two alleles in the population (Sump et al., 2022) in either SDC-His or SDC-His + 10mM 3-AT. **(C)** Average difference in Rpb1-MN between *gcn4Δ* (top) and *gcn4-pd* (bottom; mutant – *GCN4*, ΔCPMn) upstream of 246 Gcn4-target genes for cells grown + amino acids (left) or - amino acids for 1 hour (Supplementary Table 1). A region spanning 700 bp upstream and 400 bp downstream of the TSS (hashed vertical line) is displayed and the color scale reflects the difference between wild type and each mutant. **(D)** Metasite plot showing average Rpb1-MN CPMn from cells grown +amino acids (left) or - amino acids (right) for *gcn4Δ* (blue), *gcn4-pd* (purple), and *GCN4* (yellow) strains. Top: Rpb1-MN -sMNase CPMn over TATA boxes ± 250bp upstream of 173 Gcn4-dependent genes (Supplementary Table 1; Rhee and Pugh, 2012). Bottom: Rpb1-MN CPMn over 284 Gcn4 binding sites (Gcn4BS) ± 250bp upstream of 130 Gcn4 target genes (Supplementary Table 1). (**E**) The average change in Rpb1-MN by ChEC-seq2 at each gene-region in cells shifted into media lacking amino acids (SD+Uracil) vs. cells shifted into media with amino acids (SDC) is plotted (SD+Uracil – SDC, ΔCPMn) for each strain (*GCN4*, purple; *gcn4-pd*, green; *gcn4Δ*, grey). Rates *k*_2_, *k*_-2_, *k*_-3_, and *k*_4_ were re-fit to the observed Rpb1-MN occupancy at 287 Gcn4-dependent genes in wild type cells grown in the absence of amino acids and yielded 1057 STM models (blue). The rates from these best-fit models were adjusted to fit the observed changes in Rpb1-MN over each gene region in *gcn4-pd* and *gcn4Δ* strains. For *gcn4-pd*, an increase in dissociation from the promoter fit the empirical findings (τ*k*_-3_, red; Table 2) or a combined increase in promoter dissociation and decrease in transfer from UAS to promoter (τ*k*_-2_τ*k*_-3_, orange; Table 2). For *gcn4Δ*, a decrease in recruitment to UAS fit in the model the empirical findings (τ*k*_1_, black; Table 2)

ChEC-seq2 against Rpb1-MN was performed in *GCN4*, *gcn4Δ* and *gcn4-pd* mutant strains grown in the presence or absence of amino acids. This experiment identified 287 genes that showed a log_2_ fold-change (LFC) of 1 or greater (p. adj < 0.05) in the *GCN4* strain upon amino acid starvation, but not in the *gcn4Δ* strain (Table S1). These genes were strongly enriched for genes involved in amino acid metabolism (*p* = 3e-46; GO term 0006520) and strongly overlapped with Gcn4 targets (Bonferroni-adjusted *p* = 1e-10 from Fisher Exact test comparing overlap with targets defined near high-confidence Gcn4 ChEC-seq2 sites; VanBelzen et al., 2024). In the presence of amino acids, neither the *gcn4Δ* nor *gcn4-pd* mutations affected Rpb1 occupancy at the 287 Gcn4-dependent genes (Figure 7C, left column). However, upon amino acid starvation, strains lacking Gcn4 showed a stark decrease in Rpb1 recruitment upstream of the TSS that spanned both the UAS and promoter region (Figure 7C, top panel). The *gcn4-pd* mutation resulted in a more modest decrease in Rpb1 specifically over the promoter (Figure 7C, bottom). This suggested that the recruitment of RNAPII to the UAS region is dependent on Gcn4, but not on the PD_GCN4_.

Consistent with this possibility, Rpb1 cleavage adjacent to the TATA boxes near the Gcn4 target genes and over Gcn4 binding sites was strongly decreased by loss of Gcn4 (*p* < 2e-16; Kolmogorov-Smirnov test comparing the mean cleavage pattern over 173 TATAs; Figure 7D, right panels). In the *gcn4-pd* strain, Rpb1 cleavage was decreased over the TATA box (p = 4e-5; Figure 7D, top right), but not over the Gcn4 binding site (Figure 7D, bottom right). Thus, the PD_GCN4_ impacts the association of RNAPII with promoters, but does not affect RNAPII binding to the UAS.

Finally, we compared the effects of adjusting the rates of each step in our kinetic model to the effects of the Gcn4 mutations on Rpb1 occupancy (Figure 7 – Supplement). The effects of loss of Gcn4 agreed well with simply decreasing RNAPII recruitment to the UAS alone (*k*_1_, resulting in less RNAPII to move from the UAS to the promoter; Figure 7E). For the *gcn4-pd* mutant, increasing the dissociation of RNAPII from the promoter (*k*_-3_) either alone or in combination with decreasing the rate of transfer of RNAPII from the UAS to the promoter (k_2_) agreed well with these data (Figure 7E). This suggests that Gcn4 recruits RNAPII to the UAS through its activation domains and that its interaction with the NPC stabilizes promoter-bound RNAPII.

## Discussion

Understanding complex biological mechanisms requires multipronged, multidisciplinary approaches. Each approach has strengths and weaknesses but together, they provide a more complete picture. Our current understanding of RNAPII transcription, involving the dynamic collaboration of dozens of proteins, is the product of biochemical, structural, genetic, cell biological and genomic approaches. From decades of such work, we have an excellent working model for this critical biological process. Biochemical, structural and cell biological approaches (and, in some cases, genetic approaches) can be biased by the particularities of the model system(s). For this reason, global approaches provide an essential perspective to assess the generality of the conclusions from more focused studies. Our current global perspective of molecular biology is dominated by a single technique: chromatin immunoprecipitation, coupled with next generation sequencing (ChIP-seq) and its derivatives. Indeed, ChIP-seq is the sole method used to define DNA binding and chromatin state by the ENCODE and modENCODE Consortium (Landt et al., 2012). Such a methodological monoculture is problematic if there are ways in which ChIP falters in detecting important interactions (Park et al., 2013; Teytelman et al., 2013).

### ChIP vs ChEC

For proteins that bind to DNA at specific sites, ChIP-seq and ChEC-seq2 generally agree. For example, high confidence binding sites for ssTFs show excellent agreement between either ChIP-seq or ChIP-exo and ChEC-seq (Donczew et al., 2021; VanBelzen et al., 2024; Zentner et al., 2021, 2015). Likewise, mapping the associations of PIC components, Mediator or the kinases associated with transcription by ChEC-seq2 was very similar to such maps produced by ChIP-seq (Saleh et al., 2021; Wong et al., 2014). However, some exceptions have been noted, as well. ChEC-seq with both Rif1 and Sfp1 reveals biologically sensible binding sites that were not evident from ChIP-seq (Bruzzone et al., 2021).

While both ChIP-seq and ChEC-seq2 with RNAPII gives enrichment over genes that correlates with transcription, the patterns are complementary; ChIP highlights interactions with the transcribed region (reflecting paused or elongating RNAPII) and ChEC highlights interactions with the enhancer, promoter and terminator (reflecting preinitiation or terminating RNAPII). We have validated the RNAPII enrichment reported by ChEC-seq2 in five ways. First, the maps produced by two different subunits of RNAPII are highly similar (Figure 2). Second, the RNAPII ChEC signal over promoters and UAS regions correlates well with the RNAPII ChIP signal over coding sequences and with nascent transcription rates (measured by SLAM-seq; Figure 2). Third, the cleavage by either Rpb1 or Rpb3 (as well as TFIIA and TFIIE) peaks on either side of TATA boxes, which agrees well with biochemical and structural analysis of the preinitiation complex (Figure 2 & Supplementary movie). Fourth, widespread changes in transcription are captured by changes in the Rpb1 enrichment by ChEC over all gene regions (Figure 4). Fifth, depletion of TFIIB leads to loss of Rpb1 and TFIIE, as well as an increase of TFIIA, over transcriptional start sites (Figure 5). These data, strengthened by the correlations with ChEC using factors involved in initiation and elongation, argue that the patterns of RNAPII enrichment revealed by ChEC-seq2 are biologically meaningful and fit well with the literature.

Why is there a difference between RNAPII ChIP and ChEC? While ChIP captures direct protein-DNA interactions well, it is much less able to capture indirect interactions. Additional factors that may influence ChIP enrichment include the local nucleosome occupancy, the accessibility of the epitope and the relative sensitivity of different regions to shearing by sonication. Unlike ssTFs or even PIC components that bind to directly to precise genomic sites, RNAPII interacts both indirectly (through ssTFs) and directly (in the PIC and transcribing RNAPII) with different regions, each of which is associated with distinct sets of cofactors. These differences likely impact the two methods; ChEC should detect both direct and indirect interactions with DNA, whereas ChIP should strongly favor direct interactions. Likewise, ChEC will perform better in nucleosome-depleted regions while ChIP cross-linking may be enhanced by lysine-rich nucleosomes.

ChEC detects UAS-associated RNAPII observed in single molecule biochemical experiments (Baek et al., 2021; Rosen et al., 2020) that have not been observed by ChIP-seq. This is consistent with recruitment of RNAPII to ssTFs/Mediator bound to UASs. While the enhanced RNAPII ChEC signal in intergenic regions may also reflect lower nucleosome occupancy, sMNase cleavage was not enriched over UASs like Rap1 binding sites (Figure 5E). Because such sites are also occupied by Mediator (Figure 5F), this supports a Mediator-dependent mechanism of RNAPII recruitment to the UAS (Baek et al., 2021; Rosen et al., 2020). Furthermore, it is important to highlight that the RNAPII ChEC enrichment observed over promoters and UASs is consistent with that expected from known dwell times and the rate of elongation (Figure 2). The low RNAPII ChIP signal at UASs and the high signal over coding sequences could reflect both its more direct interaction with DNA and its intimate association highly cross-linkable nucleosomes during transcription (Bintu et al., 2012; Ehara et al., 2022, 2019; Kujirai et al., 2018). However, it is less clear why the RNAPII ChIP-seq signal over the promoter is so low. ChIP successfully captures enrichment of PIC components at promoters, indicating that promoter regions can be successfully enriched by ChIP. Future studies will resolve these differences.

### ChEC with elongation factors

We also present a novel method for observing the genome-wide location of the phosphorylated forms of RNAPII (Ser2p and Ser5p) using single chain antibodies (Mintbodies) tagged with MNase. ChEC-seq2 with these Mintbodies produces patterns that agree well with total RNAPII and with the kinases responsible for these modifications. Consistent with ChIP, Ser5p RNAPII is enriched in promoters and the 5’end of active genes, while Ser2p is enriched over the body and 3’ end. Inactivation of the Kin28 Ser5p kinase results in dramatic loss of RNAPII, Ser5p RNAPII and Ser2p RNAPII from active genes (Figure 5). This is consistent with an important role for Ser5p in initiation and with the observation that Ser2 phosphorylation is functionally downstream of Ser5p.

ChEC with factors involved in elongation (Ctk1, Spt5, Ser2p-RNAPII), when normalized to total RNAPII, showed greater enrichment over the CDS (Figure 3G), as expected. However, it is surprising that we also observed clear enrichment of these factors at promoters (e.g. Figure 3A, E & F). The association of elongation factors with the promoter seems to be biologically relevant. Changes in transcription correlate with changes in ChEC enrichment for these factors and modifications (Figure 4C). Blocking initiation by inhibiting TFIIH kinase led to a reduction of Ser5p RNAPII and Ser2p RNAPII over both the promoter and the transcribed region (Figure 5G). This suggests either that the true signal of these factors over transcribed regions is less evident by ChEC than by ChIP or that ChEC can reveal interactions of elongation factors at early stages of transcription that are missed by ChIP. The expectations for enrichment of elongation factors and phosphorylated CTD are largely based on ChIP data. Because ChIP fails to capture RNAPII enrichment at UASs and promoters, it is possible that ChIP also fails to capture promoter interaction of factors involved in elongation as well.

Factors important for elongation can also function at the promoter. For example, Ctk1 is required for the dissociation of basal transcription factors from RNAPII at the promoter (Ahn et al., 2009). Transcriptional induction leads to increases in Ctk1 ChEC enrichment both over the promoter and over the 3’ end of the transcribed region (Figure 4C). Dynamics of Spt4/5 association with RNAPII from *in vitro* imaging (Rosen et al., 2020) indicate that the majority of Spt4/5 binding to RNAPII does not lead to elongation; Spt4/5 frequently dissociates from DNA-bound RNAPII. Association of Spt4/5 with RNAPII may represent a slow, inefficient step in the transition to productive elongation. If so, then ChEC-seq2 may capture transient Spt4/5 interactions that occur prior to productive elongation, producing enrichment of Spt5 at the promoter.

### A role for interaction with the NPC in stabilizing the PIC

The NPC has been implicated in transcription in yeast and other organisms. In yeast, inactivation of DNA elements or transcription factors that promote interaction with the NPC leads to a quantitative defect in transcription (Ahmed et al., 2010; Brickner et al., 2012). Single molecule RNA FISH (smRNA FISH) in strains bearing mutations that blocked the interaction of the *GAL1-10* promoter with the NPC showed a decrease in the fraction of cells that exhibit transcription (Brickner et al., 2016). A mutation in the Gcn4 ssTF that blocks its ability to mediate peripheral localization and interaction with the NPC leads to a defect in expression of Gcn4 target genes (*gcn4-pd*; Figure 7; Brickner et al., 2019) and inactivation of nuclear pore proteins essential for chromatin interaction leads to a global transcriptional defect (Ge et al., 2024). Applying RNAPII ChEC-seq2, we have explored the phenotype of the *gcn4-pd* mutant. Whereas loss of Gcn4 leads to loss of RNAPII from UASs and promoters, inactivation of the PD_GCN4_ reduces the association of RNAPII with the promoter without affecting its recruitment to the UAS (Figure 7). This suggests that the PD_GCN4_ either enhances the transfer of RNAPII from the UAS to the promoter or stabilizes the association of RNAPII with the promoter. Genetic interactions between nuclear pore proteins and Mediator suggest that these two components function at the same step in transcription (Ge et al., 2024). Together with the smRNA FISH result, this suggests that nuclear pore proteins stimulate enhancer function by stabilizing RNAPII association with the PIC.

### A global model for yeast RNAPII kinetics

Because ChEC-seq2 measures global occupancy of RNAPII that includes important states that are missed by ChIP-seq, it allowed us to develop a global model for the kinetics of RNAPII transcription. Building on previous work (Rossi et al., 2021), we have modeled two classes of genes: those that show RNAPII association only with promoters (TFO) and those that show association with UASs as well (STM). For the TFO model, RNAPII is recruited directly to the promoter. For STM genes, RNAPII is recruited to the UAS and then transferred to the promoter. Subsequent steps (initiation, elongation and termination) are assumed to be the same between these two classes. Several of the rates are from the literature, while the others were fit to the experimental RNAPII enrichments over UASs, promoters, transcribed regions and 3’UTRs. While we were unable to find rates within a reasonable range of parameters that produced RNAPII occupancies matching ChIP-seq, the model identified a large ensemble of rates that produced RNAPII occupancies matching ChEC-seq2 (Figure 6B). The RNAPII occupancy from ChEC-seq2 data over highly active genes matched models that included a short dwell time over the terminator (∼ 30s), at the lower bound of what was reported in Zenklusen 2008 (mean = 56 ± 20s) and Larson et al., 2011 (mean = 70 ± 41s).

The kinetic model suggests that perturbations often have more than one effect, as expected for a dynamic, multi-step process like transcription. For example, the effects of depletion of TFIIB on RNAPII ChEC-seq2 are best modeled by both a decrease in RNAPII recruitment and an increase in non-productive dissociation of RNAPII, either from the promoter or the UAS (Figure 6C). Likewise, the effects of inhibition of Kin28 were most consistent with both a decrease in initiation and an increase in dissociation from the promoter/UAS (Figure 6D). These results suggest that the PIC is unstable and that such perturbations cause RNAPII to dissociate. This conclusion agrees with the observation that a small fraction of the polymerases that assemble at the promoter initiate transcription (Darzacq et al., 2007) and with the observation that conditional inactivation of PIC components does not preserve stable intermediates (Petrenko et al., 2019). Moreover, these results were consistent across the entire ensemble of models, showing that this is a robust effect. These models should serve as a helpful framework for future global studies of transcription.

## Methods

### Yeast strains

Yeast strains and tagging vectors used in this study are provided in Supplementary Tables S2 and S3. C-terminal MNase fusions were introduced by recombination as previously described (VanBelzen et al., 2024). Sua7 was tagged with 3xV5-IAA7 using pV5-IAA7-His3MX6, which was generated by swapping the His3MX6 marker in place of the HIS3 marker in pGZ363 (Tourigny et al., 2021). OsTir1-LEU2 was PCR amplified from pSB2271 (Miller et al., 2016) with primers that facilitated recombination at *leu2Δ0* and simultaneously restored the locus to *LEU2*. The *kin28is* mutations V21C and L83G (Rodríguez-Molina et al., 2016) were introduced by two subsequent rounds of CRISPR-Cas9 mediated mutagenesis as described (Anand et al., 2017). The *GCN4-sm* and *gcn4-pd* mutations were introduced by CRISPR-Cas9 mediated mutagenesis and are described (Ge et al., 2024).

Mintbody-MNase constructs were synthesized by Integrated DNA Technologies as gBlocks. The gBlocks were flanked by a *Hin*dIII and *Bam*HI site, which were used to clone the gBlocks into the pFA6a-NatMX6 vector (Hentges et al., 2005). The constructs were amplified from plasmid by PCR to yield amplicons flanked with homology to the *his3Δ1* locus, which were then transformed into yeast. Strains were confirmed to have the desired sequence by amplifying the modified locus from genomic DNA and sequencing. Platinum SuperFi (Thermo Fisher Scientific) was used to amplify long targets by PCR.

### Media and growth conditions

Media were prepared as described (Burke et al., 2000). Cells were grown at 30°C with shaking at 200 rpm in SDC media. YPD media was used in growth assays and in Figure 2A-C and Figure 2 – supplement 2, where cells were grown in YPD to match conditions of ChIP-seq samples. Ethanol stress was induced by growing cells in media spiked with 10% ethanol for 1 hour. Sua7-IAA7 was degraded for by treating cells with 0.5 mM Indole-3-acetic acid for 60-minutes in SLAMseq experiments or 20-minutes in ChEC-seq2 experiments. For Kin28 inhibition experiments, cells harboring the *kin28is* mutation were treated with 5 µM CMK for 60 minutes.

For SLAMseq and growth competition experiments with *GCN4-sm* and *gcn4-pd*, cells were grown in SDC and then shifted into SDC or SDC-His for 1 hour. Growth competition assays were performed as described (Sump et al., 2022) and the histidine synthesis pathway was block through the addition of 3-AT to the media. For ChEC-seq2 experiments with *GCN4-sm*, *gcn4-pd*, and *gcn4Δ*, cells were grown in YPD before shifting into either SDC or SD + uracil for 1 hour.

### ChEC-seq2

The ChEC-seq2 method was performed as described (VanBelzen et al., 2024). Cells were permeabilized and 2mM calcium was added to activate MNase activity. Reactions were stopped after genomic DNA was partially digested (see Supplementary Table S4), DNA was purified, DNA ends were repaired and ligated to an Illumina-compatible adapter (VanBelzen et al., 2024). A second adapter was incorporated through Tn5-based Tagmentation. Complete adapters and library indexes were incorporated through library amplification with Nextera XT Index Primers.

### ChIP-seq

Cell fixation and chromatin isolation was performed as previously described (Kuo and Allis, 1999) but is briefly described here for clarity. Independent cultures of BY4741 (Rpb1) and JVY022 (Rpb1-MN) were grown in YPD at 30°C, 200 rpm until cultures reached a density between 0.6 and 0.9 (OD_600_). A culture volume of 100 ml was crosslinked with 1% formaldehyde for 10-minutes at 30°C with gentle mixing. The crosslinking reaction was quenched with 0.3 M glycine for 5-minutes at 30°C with gentle mixing. A volume of 50 ml was collected by centrifugation and the pellet was washed twice in ice-cold Tris-buffered saline (20 mM Tris-HCl, pH 7.5; 150 mM NaCl), snap frozen in liquid nitrogen, and stored at −80°C for up to two weeks. Pellets were briefly thawed on ice and resuspended in 600 µl of ice-cold FA lysis buffer (50 mM HEPES-KOH, pH 7.5; 140 mM NaCl; 1 mM EDTA; 1% Triton-X 100; 0.1% sodium deoxycholate) supplemented with protease inhibitors (1 mM PMSF; 1 µg/ml Leupeptin; 1µg/ml Pepstatin A; 10 µg/ml Aprotinin). A volume of 600 µl zirconia beads (0.5 mm diameter) was added, and cells were lysed by bead-beating at 4°C in a Vortex Genie for 7 cycles of: 3-minutes on (highest setting); 1-minute on ice. The lysate was separated from the beads and brought to a final volume of 600 µl with ice-cold FA lysis buffer, which was split into two 300 µl fractions for sonication. Sonication was performed on a BioRuptor Pico (Diagenode) at 4°C for 6 cycles of: 30-seconds on (high setting); 30-seconds off. Debris was pelleted by centrifugation at 17,000 x g for 15-minutes at 4°C, and the chromatin-containing supernatant was collected.

Immunoprecipitation was adapted from (Sump et al., 2022). Dynabeads Protein G (Thermo Fisher Scientific # 10003D) were equilibrated in chilled FA lysis Buffer for 2 hours at 4°C on a rotating stand. Simultaneously, 2 mg of chromatin in a 1ml volume was incubated with 2 µl of Anti-Rpb1 (Clone 8WG16, Biolegend) at 4°C on a rotating stand. 20 µl of equilibrated Dynabeads was added to each chromatin sample and incubated overnight at 4°C on an inverting rotator. Beads were immobilized with a magnetic stand and washed four times in 1 ml of chilled Wash Buffer (50 mM HEPES-KOH, pH 7.5; 500 mM NaCl; 1 mM EDTA; 1% Triton-X 100; 0.1% sodium deoxycholate) supplemented with protease inhibitors (see above). Protein of interest was eluted in 100 µl of Elution Buffer (50 mM Tris-HCl, pH 8.0; 10 mM EDTA; 1% SDS) and crosslinks were reversed by heating overnight at 65°C. Added of 5 µl RNAse A (10 µg/µl) and heated at 37°C for 30-minutes to degrade RNA. Added 10 µl of Proteinase K (20 µg/µl) and incubated at 50°C for 1-hour. Purified DNA with QIAquick spin columns according to the manufacturer’s instructions (Qiagen # 28104). DNA fragment size was measured on a TapeStation 4150 and confirmed to be approximately 400 bp. Sequencing libraries were prepared from 0.5 ng of DNA with the MicroPlex Library Preparation Kit v3 with dual indexes (Diagenode # C05010001 and C05010004). Libraries were sequenced at NUseq on the NovaSeq X Plus (Illumina) in the paired-end, 50 bp format. Bioinformatic analysis was performed as in ChEC-seq2 (VanBelzen et al., 2024), except reads were mapped with paired-end mode of Bowtie 2 (Langmead and Salzberg, 2012) and the ChEC-specific trimming step was omitted.

### SLAMseq

SLAMseq was performed as previously described (Herzog et al., 2017) with the following modifications. Approximately 10^8^ cells were collected, resuspended in SDC-uracil + 200 µM 4-thiouracil and incubated for 6 minutes at 30°C. Cells were collected by centrifugation and frozen in liquid nitrogen. RNA was extracted from cell pellets as described (Schmitt et al., 1990), and purified with the Monarch Total RNA Miniprep Kit (New England Biolabs). Alkylated RNA was purified with the Monarch RNA Cleanup Kit (New England Biolabs). RNA quality was confirmed after each purification with a TapeStation 4150 (Agilent). Sequencing libraries were prepared from 150 ng RNA the QuantSeq 3’ mRNA-Seq Library Prep Kit (FWD) kit (Lexogen). Sequencing was performed on a HiSeq 4000 (Illumina) in the single-end, 50 bp format at the Northwestern University NUseq core facility. In the case of SLAMseq performed with JBY555 (*gcn4-pd-GFP*) and JBY556 (*GCN4sm-GFP*) (Ge et al., 2024), cells were shifted into SDC-uracil with 2 mM 4-thiouracil for 6 min.

Reads were mapped with SlamDunk (Herzog et al., 2017) to the S288C genome (build R64-3-1) and binned into genes classified as Verified or Uncharacterized by the Saccharomyces Genome Database. This yielded counts values for 5925 genes. Counts files were analyzed in R with DESeq2 (Love et al., 2014) to identify differentially expressed genes between conditions.

### Immunoblotting

Protein was isolated from cells as described (Rüegsegger et al., 2001) and quantified by BCA protein assay (#23225, Thermo Fisher Scientific). 40 µg of protein was separated on 10% surePAGE Bis-Tris gels in MOPS running buffer (#M00665, Genscript) and transferred to a nitrocellulose membrane. The membrane was blocked with 5% nonfat dry milk in TBST with 0.05% Tween 20 for 1 hour at room temperature and then probed with anti-V5 (#R960-25, Thermo Fisher Scientific) and anti-b-Actin (#GTX629630, GeneTex) primary antibodies overnight at 4°C. Membranes were washed for 5-minutes with TBST for a total of 5 washes, and then incubated with goat anti-mouse conjugated with HRP (#AP127P, Millipore-Sigma) in 5% milk TBST for 1 hour at room temperature. Washes were repeated and then HRP was activated with chemiluminescence reagents (#34075, Thermo Fisher Scientific) for 5 minutes. Blots were imaged on an c600 imaging system (Azure Biosystems).

### Computational model

We used a stochastic model to simulate the average occupancy of RNAPII along a discretized model gene (Figure 6A), assuming each step in the transcription cycle is a Poisson process. We separately modeled two classes of genes: STM genes and TFO genes. For STM genes, we assume that the association of RNAPII with the gene occurs at the UAS and is reversible, with association rate *k*_1_ and dissociation rate *k*_-1_. Next, the RNAPII transitions from the UAS to the promoter with rate *k*_2_. This rate represents an aggregate step that requires the recruitment of early general transcription factors (GTFs) such as TFIIA and TFIIB. Because these interactions are reversible, we assume RNAPII can transition back to the UAS from the promoter with rate *k*_-2_. When at the promoter, the RNAPII awaits the arrival of late GTFs such as TFIIH to form the complete PIC. This process occurs at the aggregate rate *k*_4_. While awaiting arrival of late GTFs, the RNAPII can also dissociate from the promoter with rate *k*_-3_. Once the PIC has assembled, TFIIH kinase phosphorylates the C-terminal domain of RNAPII to initiate transcription and promoter escape. This occurs with rate *k*_5_. The transcribed region is modeled as ten identical 120bp compartments, and the RNAPII moves to each succeeding compartment with rate *k*_6_. Finally, once the RNAPII reaches the terminator, it dissociates with rate *k*_7_. TFO genes are modeled similarly, with the omission of *k*_1,_ *k*_-1_, *k*_2_, and *k*_-2_, and instead introducing *k*_3_, the rate of recruitment directly to the promoter. The UAS, promoter, and terminator regions are modeled as independent 120 bp compartments. No compartment could be occupied by more than one RNAPII.

We simulated 1000 seconds of the transcription cycle to allow the system to reach steady state. We report the RNAPII occupancy of each segment of the gene over the final 60 seconds to align with the experimental procedure. The simulated data was then normalized using the *L^2^*norm and scaled to have the same magnitude as the empirical data to approximate the unit conversion to CPM or CPMn. This process was repeated across 100,000 genes and the average occupancy in each region of the gene was recorded. Simulations were performed using the Gillespie algorithm (Gillespie, 1977), a stochastic simulation method that generates statistically correct trajectories of a given system. The algorithm uses random sampling to determine the timing and sequence of state transitions that correspond to different steps in the transcription cycle. Code for the simulations is available on GitHub (https://github.com/jasonbrickner/RNAPII_kinetics_simulation).

### Model fitting

Several parameters in the model were fixed according to previously published data; *k*_1_ and *k*_-1_ were from Rosen et al., 2020; *k*_5_ was based on the residency time of TFIIH (Nguyen et al., 2021); *k*_6_ was based on an average elongation rate of 1000 bp/min (Larson et al., 2011; Zenklusen et al., 2008) and *k*_7_ is based on 56 ± 20s and 70s ± 41s (Larson et al., 2011; Zenklusen et al., 2008). Other parameters in the model were free and were fit to either ChEC-seq2 or ChIP-seq data by performing a grid search.

We evaluated each model in the grid by computing the cosine similarity between the output of the model and the empirical data. That is, we calculated the quantity

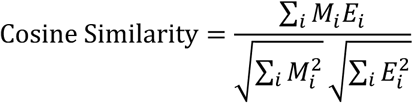

where *M_i_* is the average occupancy of the model in the *i*-th segment (UAS, promoter, transcript, or 3’UTR) and *E_i_* is the corresponding empirical data from the same segment. The cosine similarity ranges from −1 to 1, with 1 indicating perfect alignment, 0 indicating no correlation, and −1 indicating perfect inverse alignment. This measure allows us to quantitatively assess how well each model’s predictions align with the observed data simultaneously across gene regions. Rather than choosing the single model with the best fit, we elected to use an ensemble approach to more thoroughly interpret the data. In this approach, all models with cosine similarity greater than 0.995 were included in the ensemble (for ChEC-seq2). This ensemble approach allows us to explore the full space of models that are consistent with the data and avoid any spurious conclusions that may arise from the investigation of a single parameter set. The recovered ensemble of models was distributed across a manifold in parameter space, establishing required relationships between the unknown parameters (Figure 6 – Supplement 1, Figure 7 – Supplement 1). For ChIP-seq data, the model could not achieve a cosine similarity greater than 0.85, so instead we report the best fitting models to provide context. Genes with fewer than 50 nascent read counts were removed from the STM and TFO datasets, yielding 643 STM genes and 1143 TFO genes.

Based on the established functions of the proteins involved (TFIIB, Kin28 or Gcn4), we identified the rate that would be most likely influenced by the experimental perturbation and simulated the effects of perturbing that rate. If altering that rate was not sufficient to match the data, the effects of changing additional rates were explored to identify the model that best match the data. Changes to rates that did not match the empirical data are not shown. The final list of parameters used to simulate each experiment are given in Table 2.

### Data Analysis

1. Gene Classifications, Coordinates, and Regions The S288C genome sequence and annotations from build R64-3-1 were used for analysis and visualization (Engel et al., 2013). The STM and TFO gene classifications are from (Rossi et al., 2021). TATA-positions were from Rhee & Pugh, 2012. The top 150 expressed genes within each class were defined by Nascent RNA counts (SLAMseq) from the BY4741 strain grown in SDC. Similarly, expressed genes subsets were defined as genes for which there were ≥ 50 nascent RNA counts on average across 3 biological replicates. This resulted in the following number of genes per expressed subsets: STM, 643 genes; TFO, 1143 genes; TATA-containing, 597 genes. TSS and TES locations were defined by an RNA-seq dataset (Pelechano et al., 2013), when available. In cases where no TSS was available from RNA-seq, the TSS was instead taken from a CAGE-seq dataset (Lu and Lin, 2021). If neither dataset contained TSS or TES information, the median 5’UTR length (47 bp) or 3’UTR length (118 bp) was used to define these locations, respectively. Median UTR lengths were calculated from the most abundant transcript isoform for mRNAs (Pelechano et al., 2013). ChEC-seq2 signal was binned into gene regions defined as: UAS, −500 bp to −151 relative to TSS; Promoter, −150 to +25 relative to TSS; Transcript, +26 relative to TSS and −76 relative to TES; Terminator, −75 to +150 relative to TES.
2. Individual Gene Plots A region spanning 1000 bp upstream of the TSS and 1000 bp downstream of the TES is shown. Signal was smoothed with a sliding window average (window = 10, step = 5).
3. Metasite Plots Genes were aligned by TSS or TATA sequence, as indicated in the figure. 250 bp upstream and downstream of the of the aligned site was included. Signal was smoothed with a sliding window average (window = 10, step = 5).
4. Metagene Plots Metagene plots are composed of three regions: 1000 bp upstream of the TSS, the transcript (TSS to TES), and 1000 bp downstream of the TES. First, the average signal (or change in signal, where indicated) at each base pair from three biological replicates was calculated. Then, each region was divided into 100 bins and the average signal in each bin was calculated. The process was repeated for each gene, and then the average signal for each bin across all genes was calculated and is displayed in metagene plots.

## Supplementary Materials

Supplementary Movie 1: animation of the accessibility of DNA to cleavage during assembly of the closed preinitation complex, based on PDB 7nvs (Aibara et al., 2021).

Supplementary Table 1: lists of gene subsets used in this study.

Supplementary Table 2: yeast strains used in this study.

Supplementary Table 3: plasmids and oligonucleotides used in this study.

## Data Availability

Sequencing data has been deposited in the Gene Expression Omnibus at the National Center for Biotechnology Information and can be retrieved with accession numbers GSE267843 and GSE246951. Scripts used in modeling are available at https://github.com/jasonbrickner/RNAPII_kinetics_simulation.

## Funding

D.J.V. was supported by a National Science Foundation Graduate Fellowship and by T32 NIGMS GM008061. This work was supported by National Institute of General Medical Sciences grant R35GM136419 (J.H.B.).

## Acknowledgements

The authors thank Professors Vu Nguyen (University of California, San Diego), David Shore (University of Geneva), Yuan He (NU), Shelby Blythe (NU), Curt Horvath (NU) and Richard Morimoto (NU) for helpful feedback and support, members of the Brickner laboratory for helpful comments on the manuscript and Gabe Zentner for yeast strains, plasmids and technical advice.

---

**Figure 2 Supplement 1.**
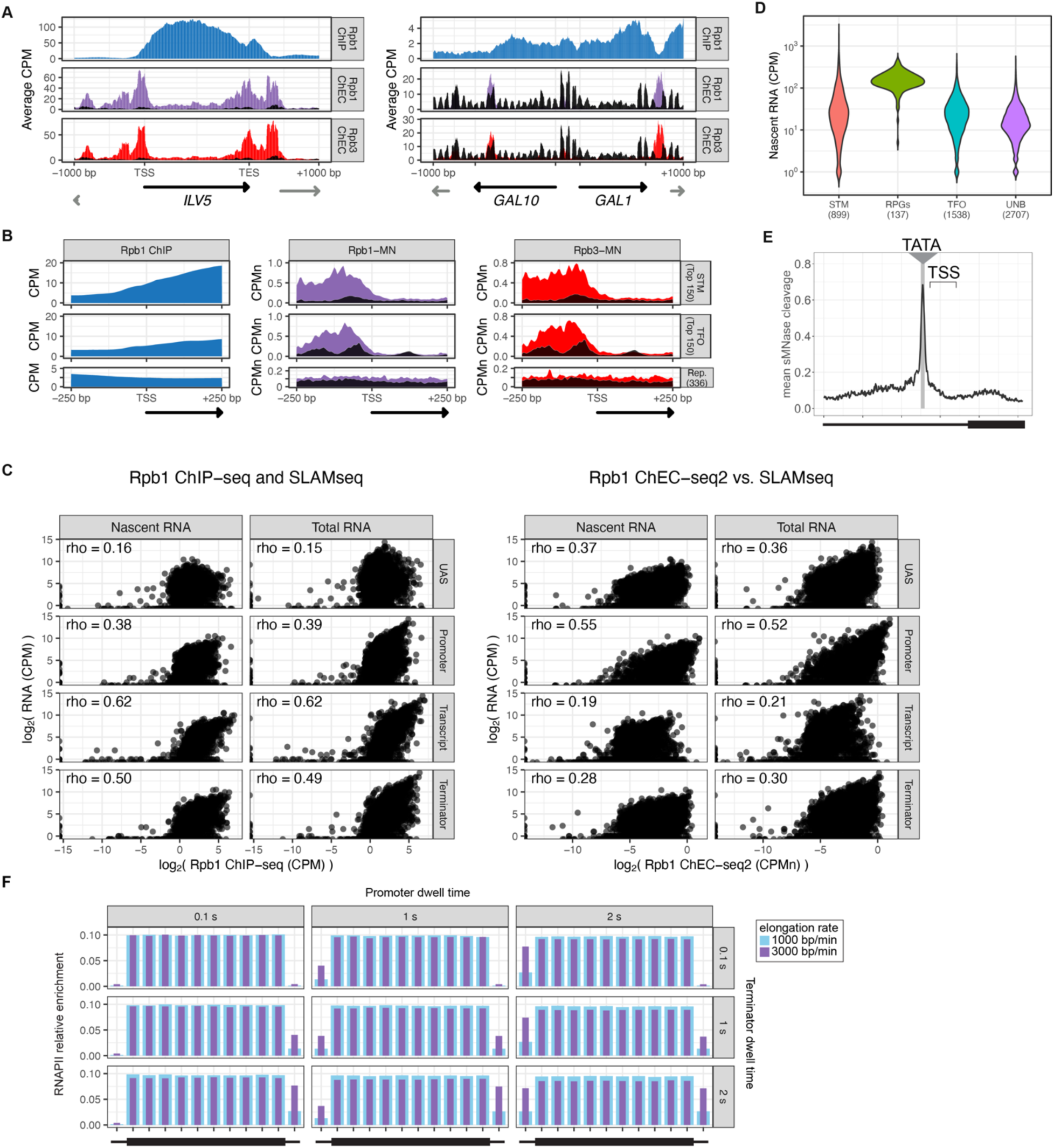
RNAPII ChEC vs ChIP. **(A)** Gene plots displaying mean counts per million reads (CPM) from Rpb1 ChIP-seq (Vijjamarri et al., 2023a) or ChEC-seq2 with Rpb1-MN or Rpb3-MN over the *ILV1* and *GAL1-10* loci. A region spanning 1 kb upstream and downstream of each locus is displayed and arrows mark the transcription start site (TSS) and transcription end site (TES). Truncated arrows represent neighboring genes that continue outside of the displayed range. Plots are smoothed with a step size of 5 and window of 10. Signal from the Soluble MNase (sMNase) control is shown in black. **(B)** Metapromoter plots showing average signal flanking the transcriptional start site ± 250 bp from 150 genes with highest expression from STM and TFO classes (Rossi et al., 2021) and 84 repressed (Rep.) genes (Supplementary Table 1). **(C)** Correlation of nascent or total mRNA levels (measured by SLAM-seq) and either ChIP-seq (left) or ChEC-seq2 (right) signal over the indicated regions of each gene. Spearman correlation coefficients for each are shown. **(D)** Nacent RNA levels from SLAM-seq for each class of genes from Rossi et al., 2021. **(E)** Metasite plot of 597 expressed, mRNA-encoding genes aligned by their TATA sequence (genes listed in Supplementary Table 1; Rhee and Pugh, 2012). Average signal from sMNase grown in rich medium is plotted. A window spanning ± 250 bp around the TATA sequence, with the TSS to the right, is shown. The location of the TATA sequence is indicated with a grey bar. The range encompassing TSSs is indicated and the rectangle below the plot designates the approximate location of the CDS. **(F)** Predicted occupancy of RNAPII based on a range of promoter dwell times (0.1 - 2 s), elongation rates (1000 - 3000 bp/min) and termination times (1 - 2 s). The transcribed region is 1200 bp divided into 10 x 120bp bins, flanked by an upstream promoter bin and downstream terminator bin. RNAPII occupancy was simulated using a minimal stochastic model. RNAPII was assumed to be immediately present at the promoter and progressed to the transcript region with a rate inverse to the promoter dwell time. It then progressed along a 1200 bp coding region with the indicated elongation rate and terminated transcription with a rate inverse to the terminator dwell time.

**Figure 2 Supplement 2.**
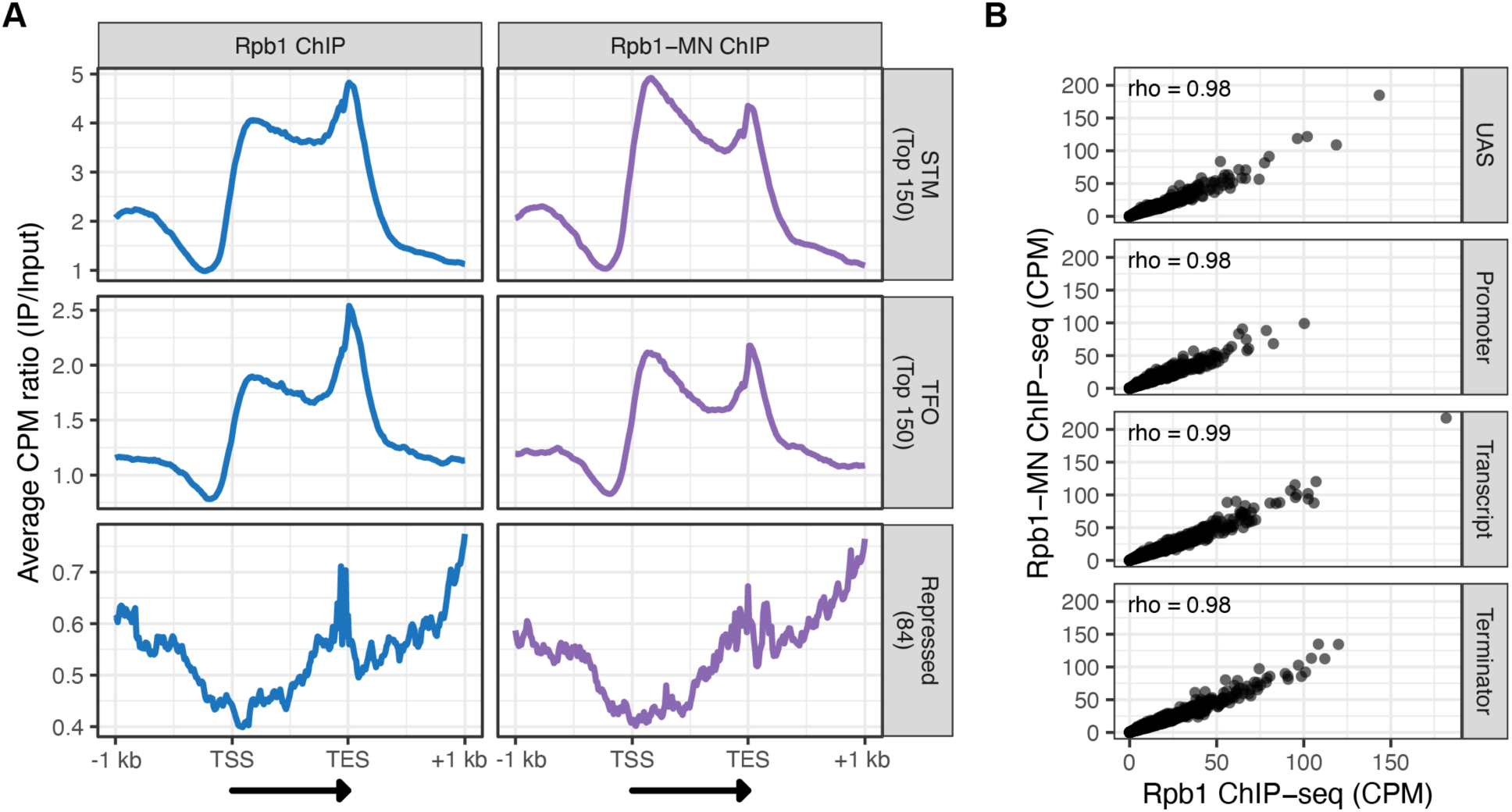
ChIP-seq against Rpb1 vs. Rpb1-MN. **(A)** Metagene plots showing the ratio of signal between IP and Input fractions (IP/Input) over subsets of genes with distinct expression levels and mechanisms of regulation. The average signal from 150 genes with highest expression from STM and TFO classes (Rossi et al., 2021) and 84 repressed genes is plotted (genes listed in Supplementary Table 1). A length-normalized transcript (arrow), 1 kb upstream of the TSS, and 1 kb downstream of the TES is shown. Rpb1 ChIP-seq (left; blue), Rpb1-MN ChIP-seq (right; purple). **(B)** Correlation of ChIP-seq against Rpb1 vs. Rpb1-MN. Signal (CPM) over the indicated regions of each gene are compared. Spearman correlation coefficients for each are shown. The average of three biological replicates is shown in (A) and (B).

**Figure 3 Figure supplement 1.**
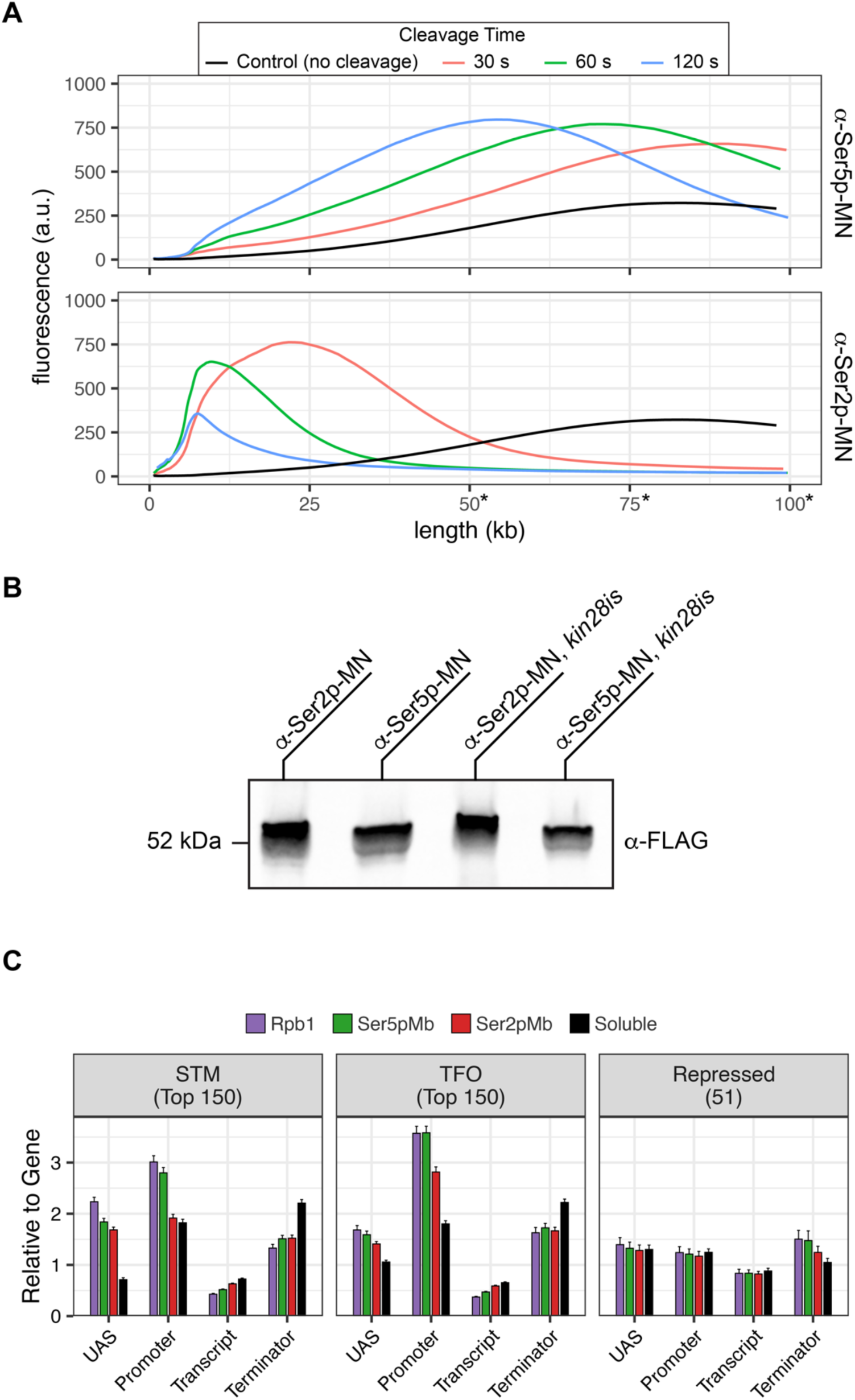
Mintbody-directed ChEC. **(A)** Genomic DNA isolated from strains expressing Ser5p-MN (JVY305) and Ser2p-MN (JVY302) was analyzed on a TapeStation 4150. MNase was activated and cleavage proceeded for 30 seconds (red), 60 seconds (green), or 120 seconds (blue). Genomic DNA isolated from cells where no cleavage occurred is shown in black. Note: absolute determination of molecular weight above 50 kb is not possible with this assay and is shown here to highlight relative changes in molecular weight between samples. **(B)** Chemiluminescent western blot of strains expressing Mintbody-MNase constructs specific to Ser2 phosphorylation (a-Ser2p-MN, JVY302) or Ser5 phosphorylation (α-Ser5p-MN, JVY305) of the CTD of RNAPII. Strains expressing each construct on the *kin28is* background are also shown (α-Ser2p-MN, JVY314; α-Ser5p-MN, JVY317). **(C)** The relative enrichment at UAS, promoter, transcript, and 3’UTR regions was calculated and normalized by region length for each gene. The average from all genes in each group is plotted. Error bars represent the standard error of the mean between three biological replicates.

**Figure 5 Supplement 1.**
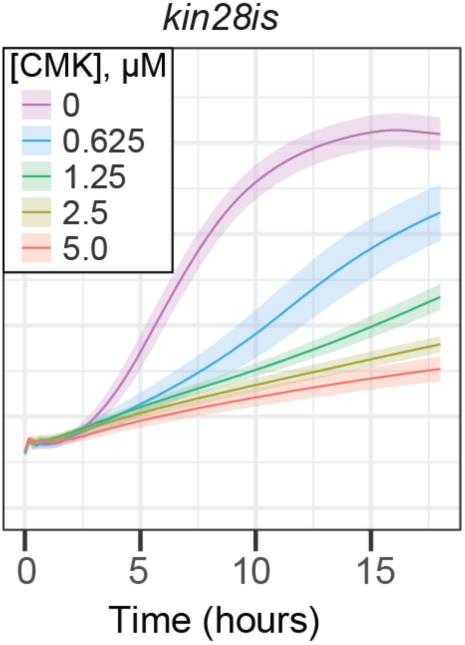
Growth eRect of CMK treatment in wild type and *kin28is* cells. **(A)** OD_600_ of *kin28is* strain grown at 30°C in synthetic complete medium with the indicated concentrations of CMK. The average ± standard deviation is plotted.

**Figure 6 Figure supplement 1.**
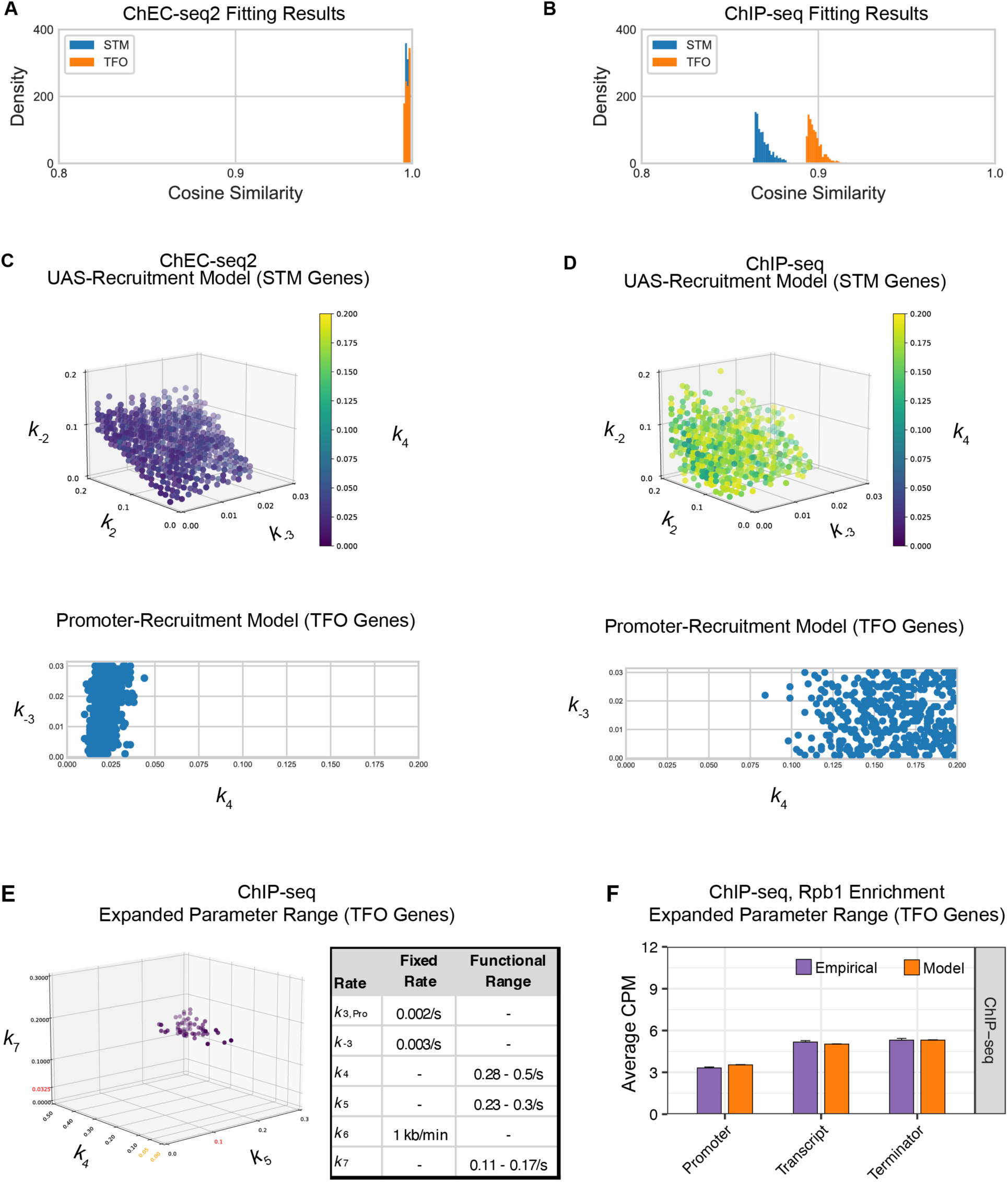
Parameter fitting of unknown transcription rates. Rates with no known value were fit to RNAPII occupancies from either ChEC-seq2 or ChIP-seq data using a grid search (see Methods). **(A-C)** For STM model, rates *k*_2_, *k*_-2_, and *k*_4_ were explored in the range [0, 0.2] and *k*_-3_ in the range [0, 0.03]. For TFO model, only rates *k*_-3_, and *k*_4_ were fit, in the same range. **(A)** Distribution of cosine similarity for the model ensemble when fit to ChEC-seq2 data. Cosine similarity of 1 indicates perfect alignment, 0 indicates no correlation, and −1 indicates perfect inverse alignment. **(B)** Distribution of cosine similarity for the model ensemble when fit to ChIP-seq data. **(C)** Rate combinations that fit the empirical data ChEC-seq2 data (Rpb1-MN). This resulted in 789 rate combinations for the STM model and 371 rate combinations for the TFO model. **(D)** No rate combinations resulted in a satisfactory fit to the empirical ChIP-seq data (Rpb1). Instead, an equal number of rate combinations (best-fit) as shown in (**C**) is displayed. **(E)** In an attempt to identify rates that fit the RNAPII enrichment seen by ChIP-seq, we used the promotor-recruitment model (TFO genes) and loosened previously fixed rates *k*_5_ and *k*_7_ and expanded the search range for *k*_4_ while fixing *k_-3_*. Published rates for *k_5_* and *k_7_* are displayed in red on the axes. Range from parameter fit *k*_4_ from ChEC-seq2 data (C) is shown in orange on the *k*_4_ axis. The range of values for *k_4_, k_5_*, and *k_7_* that fit the ChIP-seq data are shown in the table (Functional Range). In the idealized case *k_-3_* = 0 and *k_4_* is instantaneous then *k_7_* should be equal to the product of *k_6_* and the ratio between the average occupancy given by ChIP in the coding region and terminator of the gene (approximately 0.14 s^−1^), and *k_5_* should be equal to the product of *k_6_* and the ratio between the average occupancy given by ChIP in the coding region and the terminator of the gene (approximately 0.2 s^−1^). The functional ranges shown agree with this, as rate *k_5_* is bounded below by the idealized approximation, and rate *k_7_* is centered around its idealized approximation. **(F)** The average Rpb1 signal (purple) from ChIP-seq over the indicated regions from TFO-class genes that are expressed in SDC. RNAPII enrichment resulting from rate combinations shown in (E) were modeled in combination with fixed rates from the literature shown in Table 1. UAS, Promoter, and 3’UTR were represented by a single 120 bp bin and the transcript region was composed of 10 sequential bins to represent a 1200 bp transcript. The average predicted occupancy for RNAPII over each region from the models (*i.e.* sets of rates) that best matched the empirical data are shown (see Methods). For Rpb1 ChIP-seq, 55 rate-combinations from the promoter model fit the empirical data from TFO-class genes. Empirical and model outcomes were compared for each gene region with a Student’s t-test, which reported no significant differences (p > 0.05).

**Figure 7 Figure supplement 1.**
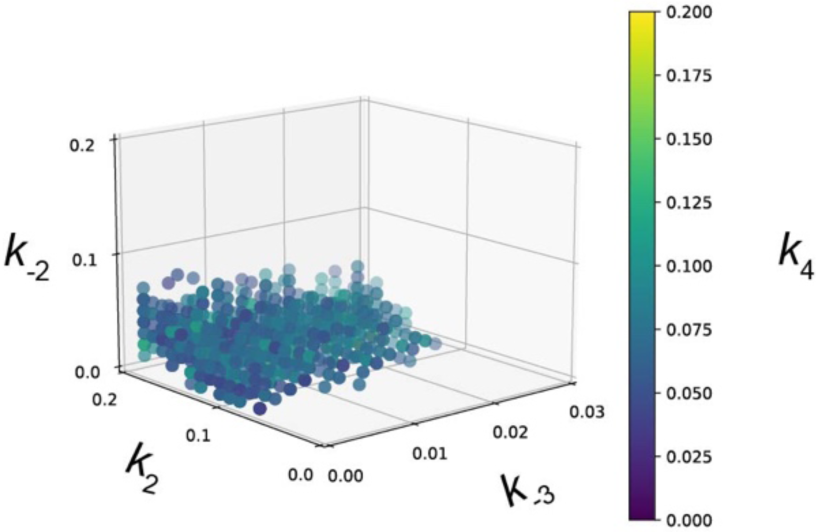
Parameter fitting of unknown transcription rates in UAS-recruitment model for Gcn4 target genes. Rates with no known value were parameter fit using a grid search (see Methods). We used the UAS-recruitment model and explored rates *k*_2_, *k*_-2_, and *k*_4_ in the range [0, 0.2] and *k*_-3_ in the range [0, 0.03]. Rate combinations that fit the Rpb1-MN ChEC-seq2 data from 287 Gcn4-target genes under amino acid starvation conditions. The fitting procedure resulted in 1057 rate combinations that fit the empirical data.

## Notes

### Competing Interest Statement

The authors have declared no competing interest.

### Summary of Updates

This version has been revised to address reviewers' comments for publication in eLife as follows: We have changed the title of the manuscript to Chromatin endogenous cleavage provides a global view of yeast RNA polymerase II transcription kinetics. Text 1. Additional discussion of the patterns for elongation factors added. 2. Small text changes throughout. 3. Added to Methods: ChIP-seq performed to confirm that the MNase fusion proteins are able to produce the expected pattern for ChIP. Figures 1. Updated legend-image in Figure 2F to reflect correct colors 2. Added Figure 2 supplement 1F: RNAPII enrichment with shorter promoter dwell times 3. Added Figure 2 supplement 2 with ChIP-seq outcomes (and text legend) 4. Removed gene numbers in Figure 5C and put them in the legend. 5. Substituted Med1 and Med8 ChEC over Rap1 sites in Figure 5F. 6. Moved kin28-is growth inhibition to Figure 5 Supplement 1. 7. Substituted a new panel overlaying the RNAPII enrichment over UASs or promoters for all three strains in Figure 7D. 8. Improved the labeling and legend of Figure 7E.

